# Myosin Va Transport of Liposomes in Three-Dimensional Actin Networks is Modulated by Actin Filament Density, Position, and Polarity

**DOI:** 10.1101/526640

**Authors:** Andrew T. Lombardo, Shane R. Nelson, Guy G. Kennedy, Kathleen M. Trybus, Sam Walcott, David M. Warshaw

## Abstract

The cell’s dense three-dimensional (3D) actin filament network presents numerous challenges to vesicular transport by teams of myosin Va (MyoVa) molecular motors. These teams must navigate their cargo through diverse actin structures ranging from Arp2/3-branched lamellipodial networks to the dense, unbranched cortical networks. To define how actin filament network organization affects MyoVa cargo transport, we created two different 3D actin networks *in vitro*. One network was comprised of randomly oriented, unbranched actin filaments; the other was comprised of Arp2/3-branched actin filaments, which effectively polarized the network by aligning the actin filament plus-ends. Within both networks, we defined each actin filament’s 3D spatial position, using STORM microscopy, and its polarity by observing the movement of single fluorescent, reporter MyoVa. We then characterized the 3D trajectories of fluorescent, 350 nm fluid-like, liposomes transported by MyoVa teams (~10 motors) moving within each of the two networks. Compared to the unbranched network, we observed more liposomes with directed and fewer with stationary motion on the Arp2/3-branched network. This suggests that the modes of liposome transport by MyoVa motors are influenced by changes in the local actin filament polarity alignment within the network. This mechanism was supported by an *in silico* 3D model that provides a broader platform to understand how cellular regulation of the actin cytoskeletal architecture may fine-tune MyoVa-based intracellular cargo transport.

**SIGNIFICANCE STATEMENT:** Intracellular transport of critical cellular components (e.g. vesicles, organelles, mRNA, chromosomes) is accomplished by myosin Va molecular motors along complicated three-dimensional (3D) networks of actin filaments. Disruption of these transport processes leads to debilitating human disease (e.g. Griscelli Syndrome), while rearrangement of the 3D actin cytoskeleton is a hallmark of malignant cancers. We found that the various modes of motion (stationary, diffusive-like or directed) describing how teams of myosin Va transport 350nm liposome cargos are determined by the 3D position and polarity of the actin filaments within the network that the myosin Va motors interact with. This study demonstrates that the 3D actin filament organization within the network can serve as a potent regulator of myosin Va motor-based intracellular transport.

## INTRODUCTION

To maintain homeostasis through critical cellular functions, cells are required to continuously transport cargo (e.g. vesicles, organelles, mRNA, chromosomes) through the crowded intracellular environment. This task is accomplished by molecular motors; biological machines that transport a variety of intracellular cargo along the cell’s labyrinth of cytoskeletal filaments (Hirokawa, 1998; Mehta et al., 1999; Ross et al., 2008; Vale et al., 1996). Cargos targeted for secretion at the membrane rely on kinesin and dynein motors for long range (~10μm) transport on microtubules. Final delivery from the cell cortex to the membrane requires shorter range transport (1-3μm) by myosin motors on actin filaments (Fig.1A) (Hendricks et al., 2010; Kapitein et al., 2013; Langford, 2002). One such myosin motor, myosin Va (MyoVa), is responsible for the transport and anchoring of lipid-bound vesicle cargo in a wide array of cell types (e.g. melanosomes, chromaffin cells, neurons) (Balasanyan & Arnold, 2014; Rosé et al., 2003; Wu, Bowers, Rao, Wei, & Hammer III, 1998).

**Figure 1.**
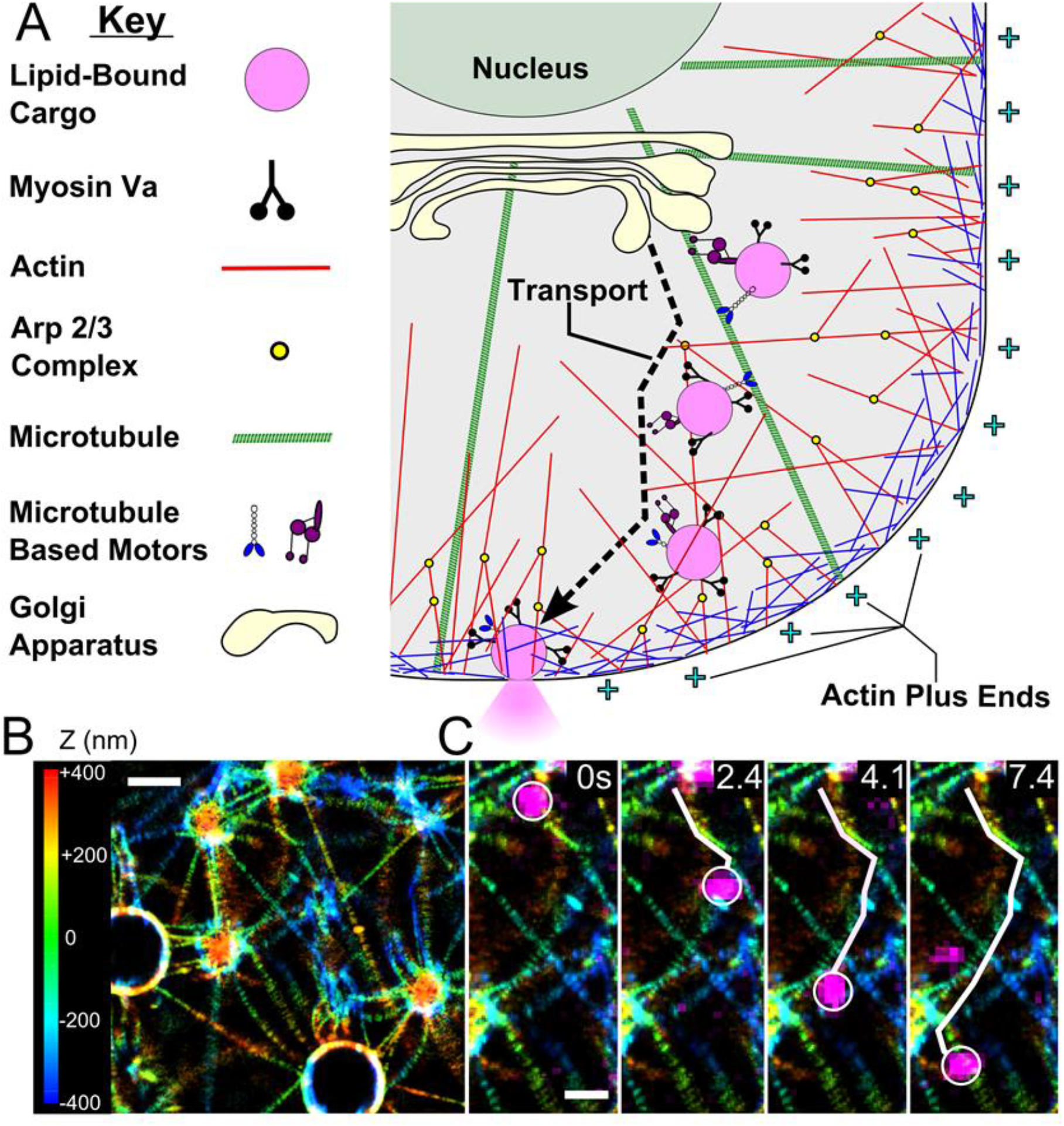
Myosin Va Teams Transport Intracellular Cargo Through Networks of Actin. **(A)** Lipid-bound cargo is produced and packaged at the interior of the cell within the Golgi. These cargos are first transported along microtubule tracks followed by hand-off to MyoVa for distribution and final delivery to sites of secretion at the cell membrane. **(B)** Super resolution, 3D STORM reconstruction of an *in vitro* actin network. Actin filaments are strung between silica beads of varying diameters, which support the network and maintain a 3D organization. Color represents Z-position. Scale 2μm. **(C)** Overlay of 350 nm vesicle trajectory (magenta) by teams of MyoVa within a 3D actin filament network (colored by Z-position). Scale 1μm.

Cells and their actin cytoskeletal networks are three-dimensional (3D). In these networks, each actin filament is polarized by the structural arrangement of the actin monomers in the filament to create a plus- and minus-end. MyoVa detects this polarity and walks exclusively towards the plus-end. However, the 3D position and the polarity alignment of the actin filaments can vary widely within a cell as a cargo vesicle is transported through and along different actin cytoskeletal elements (Blanchoin, et al., 2014; Pollard, Blanchoin, & Mullins, 2000; Svitkina et al., 1997). How cells employ teams of MyoVa motors to navigate their cargo through such varied cellular actin networks in order to ensure proper cargo delivery to its destination is far from certain. The importance of this question is highlighted by genetic mutations of MyoVa in humans, which result in devastating neurological diseases (Pastural et al. 1997). The expression of a dominant negative MyoVa construct in cultured cells suggests that these diseases result from failure to properly transport intracellular cargo, leading to cargo aggregation at the cell’s interior (Gross et al., 2002). While cellular studies provide context for MyoVa’s multiple roles *in vivo*, the cell’s complexity makes interpretation of MyoVa’s specific contributions to transport difficult. Thus, complementary *in vitro* work using simplified cargo transport by molecular motors on cytoskeletal model structures (Ali et al., 2007; Hariadi et al., 2014; Nelson et al., 2014; Ross et al., 2008; Yildiz et al., 2003) may provide a mechanistic basis underlying intracellular cargo transport. However, these *in vitro* studies generally rely on rigid, non-physiological cargo (e.g. quantum dots, beads, DNA origami) and lack the 3D nature of the cell’s cytoskeletal filament network (e.g. through the use of glass-surface-bound filaments).

More recently, investigators have begun to build complexity both *in vitro*, and in cellular model systems, by exploring the transport of cargos in 3D (Bálint et al., 2013; Bergman et al., 2018; Lombardo et al., 2017; Verdeny-Vilanova et al., 2017). When compared to 2D model actin networks bound to glass surfaces, MyoVa motors can move cargo greater distances, and can maneuver cargo past physical obstructions when actin filaments are suspended in 3D (Ali et al., 2002; Lombardo et al., 2017). This result suggests that the 3D position and polarity of actin filaments can influence the navigation of cargo by MyoVa. However, dense actin networks in cells present a seemingly insurmountable obstacle to transport, as motors on a single cargo may contact several (2-6) filaments simultaneously (Snider et al., 2004); setting up a potential “tug of war” between engaged motor teams on the cargo surface and thus resulting in a stationary or tethered cargo. Through the alignment of actin filament plus-ends within the network (i.e. introducing a polarity bias) (Ross et al., 2008; Svitkina et al., 1997), directed MyoVa cargo transport may then be possible (Hariadi et al., 2014). Therefore, the local 3D position and polarity of actin filaments in a network may be tunable regulators of MyoVa transport.

To investigate how local actin position and polarity affect MyoVa transport, we adapted our previously reported 3D actin model system (Lombardo et al., 2017) to create 3μm thick, 3D actin filament networks of similar filament density to those found in cells (Fig. 1B) (Snider et al., 2004). Using 3D super-resolution Stochastic Optical Reconstruction Microscopy (STORM), we imaged and determined each actin filament’s 3D position within the network and subsequently defined the polarity of each filament using a fluorescent reporter technique (Huang et al., 2008; Tas et al., 2017). The local polarity of filaments within the network was manipulated by nucleating branched filaments, using the Verprolin, Cofilin, Acidic (VCA) domain of Wiskcott-Aldrich Syndrome Protein (WASP) and Actin Related Protein 2/3 (Arp2/3) complex (Pollard et al., 2000; Reymann et al., 2012). This created Arp2/3-branched filaments where the plus-ends of the nucleated branches face in a similar direction as the mother filament. Using physiologically relevant, liposome cargos with constitutively active MyoVa motors on their surface, motor-cargo complexes were challenged to navigate these 3D actin networks (Fig. 1B,C). The increase in local plus-end actin filament alignment in the Arp2/3-branched network biased motor-cargo complex transport towards directed motion, as compared to networks of unbranched filaments, where the actin filament polarity was arranged randomly. To explain this result, we used an *in silico*, 3D mechanistic model that characterizes the contribution of every actin filament-engaged motor on the fluid-like liposome surface to define the liposome’s transport behavior (Lombardo et al. 2007). This model recapitulates the experimental observations and demonstrates that different modes of liposome motion arise from motors either producing cooperative forces aligned in a similar direction, which favors directed motion, or engaging in a tug-of-war, which favors stationary/tethered liposomes. Thus, the local actin filament polarity within dense actin networks of the cell can modulate the modes of motion for MyoVa-bound cargo.

## RESULTS

### Actin network characterization

A 3μm thick network of randomly oriented actin filaments was created off the glass surface by suspending actin filaments between poly-L-lysine-coated silica beads of diameters ranging from 500 nm to 3000 nm within a microfluidic chamber (Fig. 1B). The mixture of bead diameters prevented network compaction, with actin filaments spanning 2.3 ± 0.7μm (mean ± SEM) between beads and creating numerous 3D actin filament intersections (Fig. 1B), thereby mimicking cellular actin filament networks (Kapitein et al., 2013). Alexa-647-labeled phalloidin was used to label the actin, which was then imaged using super-resolution 3D STORM by incorporating an intentionally induced astigmatism into the light path (Huang et al., 2008; Lombardo et al., 2017). A 3D linear fit to STORM localization data for fluorophores along the filament was used to identify the spatial relationship of each individual actin filament within the network (Fig. 2), with a precision of ±6.5 nm, based on the Root Mean Squared Deviation (RMSD) for the fit. We designed two separate actin networks to emulate the types of cellular networks where MyoVa is known to play a critical role in cargo transport: (1) unbranched cortical actin filament networks (Rosé et al., 2003); (2) Arp2/3-branched networks originating from the Golgi apparatus in many secretory cells (Chen, Lacomis, Erdjument-Bromage, Tempst, & Stamnes, 2004; Matas, Martínez-Menárguez, & Egea, 2004), at the leading edge of motile cells (Pollard & Cooper, 2009), and in dendritic spines of neurons (Wagner et al., 2011). To create branched networks, we polymerized actin filaments in the presence of Arp2/3, which created filaments with multiple branch points (Fig. 2C inset), and then incorporated these into our 3D suspended networks (Fig. 2C).

**Figure 2.**
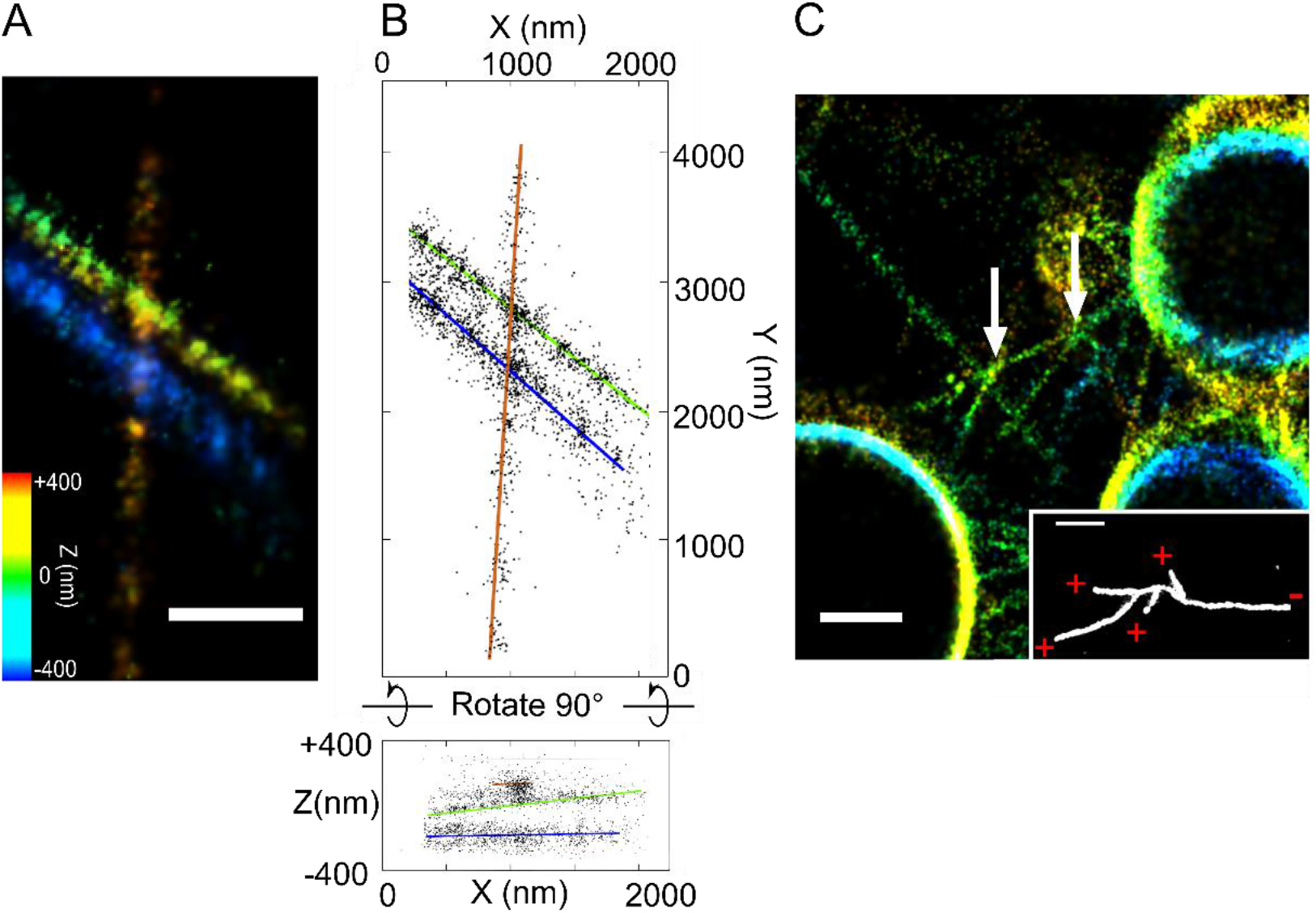
Measurement of Actin Filament Position Within a 3D Network and Creation of Arp2/3-Branched Networks. **(A)** 3D STORM reconstruction of three actin filaments with Z-position indicated by color. (Scale: 1μm) **(B)** Fits through the STORM localization data of the identical actin filaments shown in (A), identify each filament’s 3D position. Error on the actin orientation precision was measured between the fit and the STORM data points for each filament (RMSD; Orange filament: 6.9 nm n=435; Green filament: 6.3 nm n=695; Blue filament: 6.3 nm n=465). Rotation of the localization data and fits by 90 degrees reveals the 3D orientation of the filaments. **(C)** A 3D network of branched filaments (white arrows) was created using filaments polymerized with Arp2/3. (Scale: 1 μm) Inset: Prior to introduction of the Apr2/3-branched filaments into the 3D network, a small sample was separated from the polymerized stock, diluted and then imaged using STORM to confirm the presence of branched filaments. (Scale: 1 μm).

### Modes of MyoVa-dependent liposome motion are related to the type of actin filament network

The 3D trajectories of 350±32 nm (mean ± s.d.) synthetic, lipid membrane liposomes with ~10 MyoVa bound were tracked with high spatial (XY, 18 nm; Z, 30 nm) and temporal (50 ms) precision (Lombardo et al., 2017), as they navigated actin filament networks (see Methods). The average number of liposome-bound motors was determined using fluorescence photobleaching; while liposome diameters were measured by dynamic light scattering (Lombardo et al., 2017; Nayak & Rutenberg, 2011; Nelson et al., 2014). The use of DOPC phospholipid liposome membranes, which are fluid at room temperature, allowed for free diffusion of motors across the liposome surface (Nelson et al., 2014). When liposomes were transported on a single, isolated suspended filament, they were limited only by the length of the actin and moved with a velocity of 389±163 nm/s (mean ± SEM), which matches values previously reported (Lombardo et al., 2017). However, when MyoVa-driven liposomes were tracked within an unbranched filament network (Fig. 1C), the behavior of the cargo became more complex, often switching between stationary, diffusive-like, and directed motions (Fig. 3A). To describe these three modes of motion, we used Mean Squared Displacement (MSD)-based segmentation analysis to calculate the diffusive exponent, or alpha value (α), which is a measure of the liposome’s mode of motion (Nelson et al., 2009; Heaslip et al., 2014). Specifically, the motion modes were defined by ranges of alpha values as follows: stationary (α≤0.67), diffusive-like (0.67 < α < 1.33), or directed (α ≥ 1.33) (Fig. 3A) (Heaslip et al., 2014; Weihs, Mason, & Teitell, 2007) (see Methods). The cutoff for the stationary mode of motion was determined by tracking the positions of MyoVa-liposomes bound to actin at 0 nM ATP conditions (rigor) (Fig. S1). Similarly, the cutoff for directed motion was determined by tracking the position of liposomes transported along single actin filaments at 1mM ATP (saturating) conditions, which exhibit long, robust transport (Fig. S1) (Nelson et al., 2014). These alpha value cutoffs are similar to those used for *in vivo* transport modes of insulin granules and early endosome cargos (Heaslip et al., 2014; Zajac, Goldman, Holzbaur, & Ostap, 2013).

**Figure 3.**
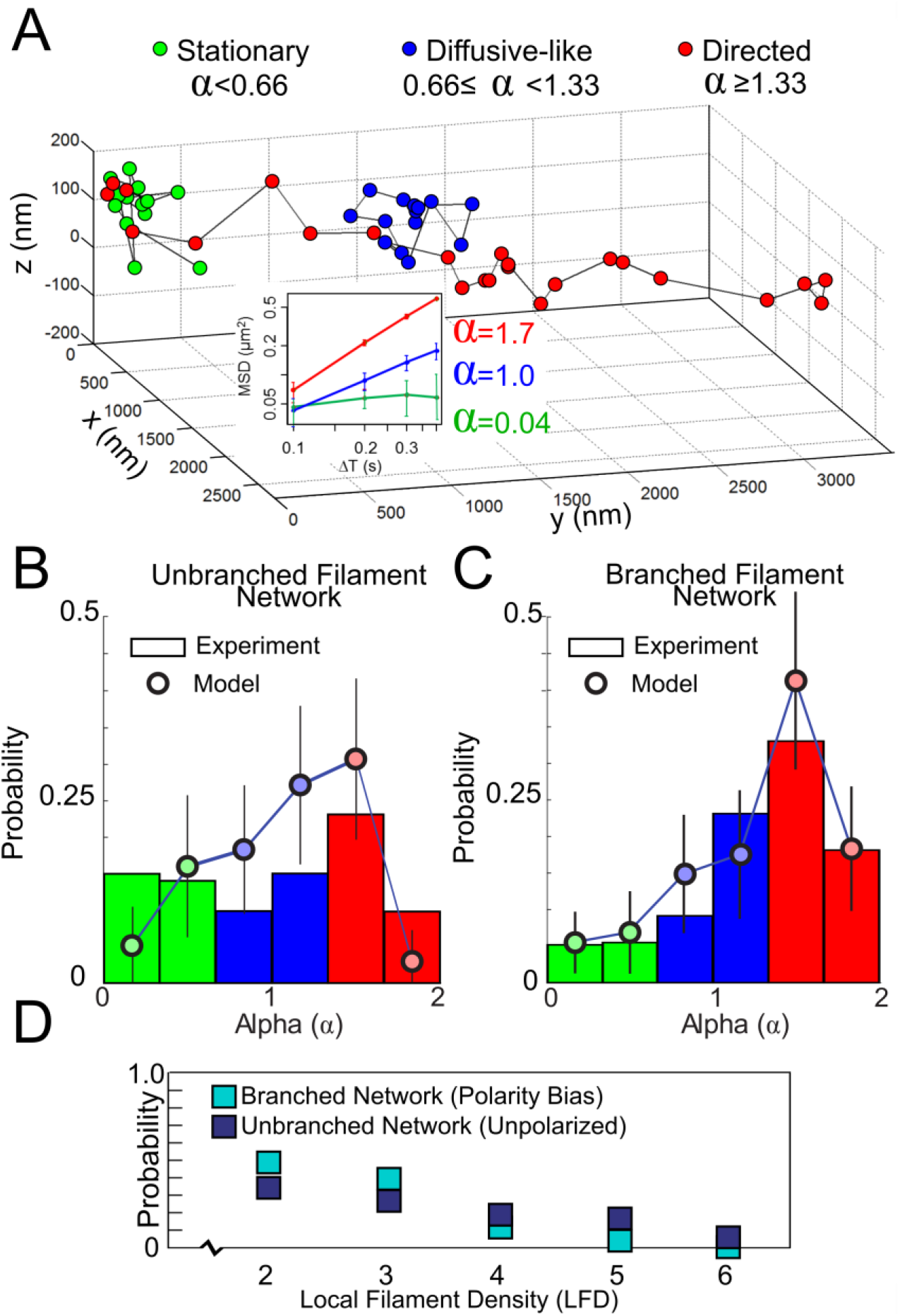
Tracking Liposomes in 3D and MSD Modes of Motion Analysis. **(A)** Representative example of the 3D trajectory of a 350 nm liposome transported within an unbranched actin filament network, which exhibits multiple modes of motion as indicated by color (Directed=Red; Diffusive-like=Blue; Stationary=Green). The liposome’s MSD for the segmented regions are plotted vs. time (ΔT) on a log-log axis (inset). The slope of the linear fit to the plotted data represents the diffusive coefficient, alpha (α) reveling the three separate modes of motion within the single trajectory: Stationary (α=0.04), diffusive (α=1.0), and directed (α=1.7). **(B)** Probability distributions of α values in 3D unbranched networks (n=329 liposomes) and **(C)** in Arp2/3-branched filament networks (n=245). Probability distributions are colored by their alpha value to indicate modes of motion as defined in (A). Mechanistic *in silico* model results (circles) overlaid on top of experimental results. **(D)** Probability distribution plot for the Local Filament Density (LFD) between Arp2/3-branched (cyan) and unbranched (Navy blue) 3D networks of filaments. Liposomes moving on a single filament were considered not in a network and excluded. No significant difference was found between the distributions (p=0.69).

We then compared the distribution of alpha values for liposomes moving within a network of actin filaments (i.e. in contact with 2-6 filaments) that was made from either simple, unbranched filaments, or Arp2/3-branched filaments (Fig. 3B,C). Liposomes moving along single filaments, as well as those undergoing diffusive-like motion, which were not in proximity (>700 nm) of any filaments, were considered not in contact with a network and were excluded from the dataset. Within unbranched filament networks, the liposome modes of motion were approximately equally distributed between directed (38%), diffusive-like (29%), and stationary (33%) (n=329) (Fig. 3B). Strikingly, within Arp2/3-branched 3D networks, the liposome modes of motion showed a statistically significant shift towards more directed motion (p=0.007) with 54% being directed, 35% diffusive-like, and only 11% being stationary (n=245) (Fig. 3C). The lifetime for stationary liposomes (~4s) was the same for Arp2/3-branched and unbranched networks (p=0.39; Table S1), suggesting that the mechanisms behind the stationary mode of motion are the same between the two networks. Therefore, the differences in the proportions of stationary liposomes in the unbranched and Arp2/3-branched networks may simply be due to the frequency of encountering local regions of the network where actin filament positions and polarities promote a stationary mode of motion (Fig. 3B,C).

### Arp2/3-branched filament networks have an increased actin polarity alignment

We hypothesized that the increase in directed and decrease in stationary modes of motion for liposomes in the Arp2/3-branched networks (Fig. 3C) resulted from either a decrease in the local actin filament density, or an increase in the alignment of the plus-ends of actin filaments, or both. To test these hypotheses, we first determined the Local Filament Density (LFD), defined as the number of filaments a liposome could potentially contact (Fig. 3D). LFD was determined by calculating a spherical volume, centered around the position of the liposome’s measured center of mass, that extended beyond the liposome perimeter by the reach of motors on the liposome surface (i.e. 50 nm) (see Methods). The number of filaments within this volume (i.e. LFD) was identified for every frame in a liposome’s trajectory. Liposomes transported within both Arp2/3-branched and unbranched networks showed an LFD of 2 to 3 filaments for the majority of recorded frames (Fig. 3D), with no significant difference in LFD (p=0.69) found between the two networks (Fig 3D). Therefore, differences in mode of liposome motion between the two networks were not due to differences in the network’s local actin filament density.

To determine whether differences in actin filament polarity alignment between the two networks may be responsible for the differences in the modes of liposome motion (Figs. 3B,C), we determined the polarity of each actin filament in contact with a liposome. By employing a myosin-cargo reporter complex, comprised of a small, ~50 nm fluorescent liposome cargo with only a single MyoVa motor (Fig. 4A) (see Methods) that processes exclusively towards the actin filament plus-end, the 3D trajectory of this reporter mapped out the polarity of each filament it moved on (Fig. 4B-D, Movie S1) (Tas et al., 2017). This polarity mapping technique was applied after 350nm liposome transport was observed within a network.

**Figure 4.**
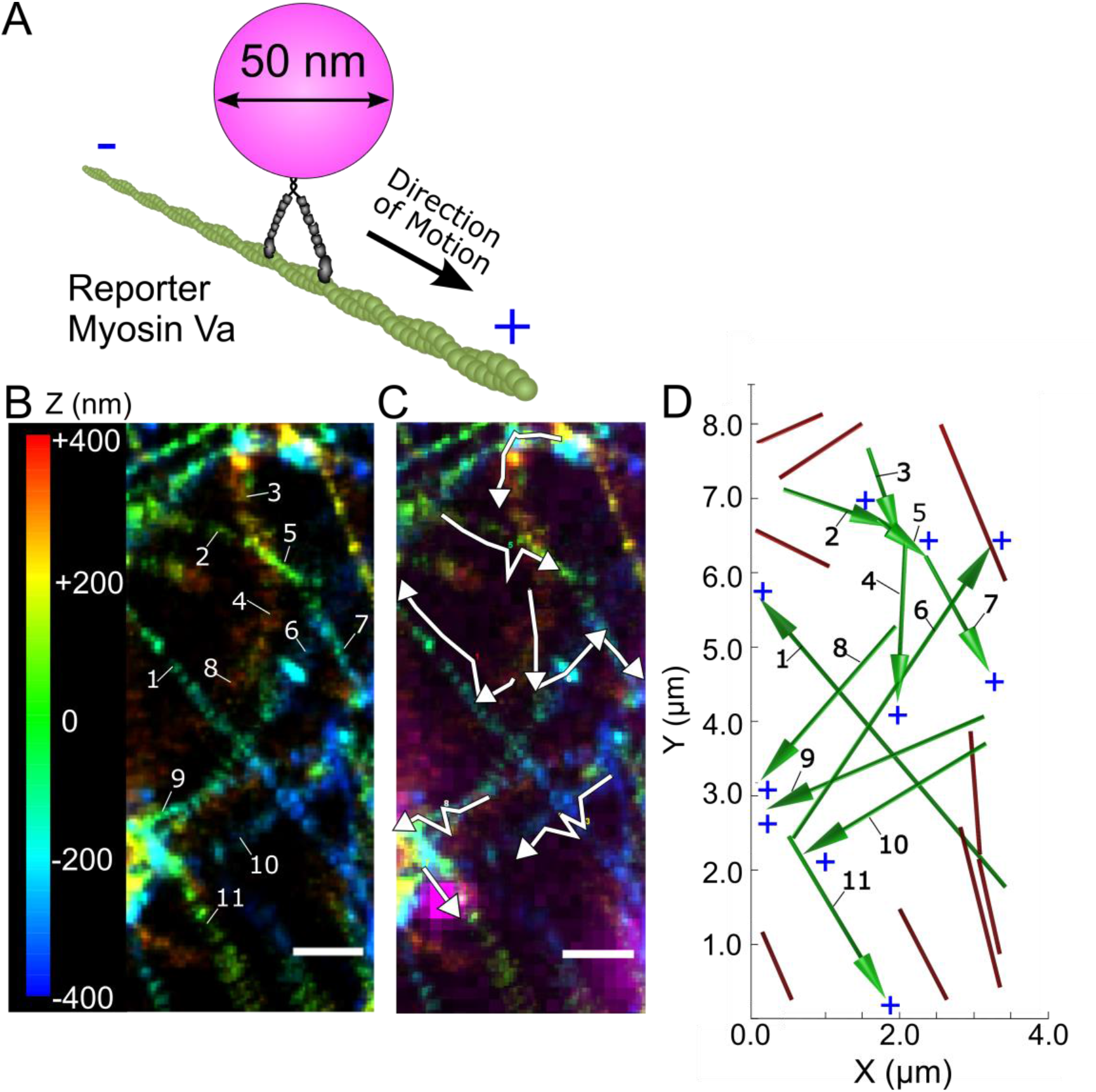
Determination of Actin Filament Polarity. **(A)** Single MyoVa were attached to 50 nm fluorescent liposome cargo to identify the polarity of filaments within a 3D network. **(B)** Actin filament network from Figure 1B,C. **(C)** Tracks of the reporter cargo (white) reveal the polarity of the actin filaments identified for the same actin network shown in (B) and Figure 1C. (Scale: 1 μm) **(D)** Combining the actin filament fits for this network with the filament polarity information determined in (C) allows for the position and polarity of every individual filament to be defined. Filaments that potentially contacted, i.e. are within 700 nm (two 350 nm liposome diameters) of the trajectory shown in Figure 1C, are colored green with the plus-end of the actin indicated by the arrow head.

Knowing each actin filament’s polarity helped define the overall Local Polarity Alignment (LPA) of the filament network in which the liposome was being transported (Fig. 5A). To begin, the polarity of an actin filament was treated as a unit vector, with the vector pointing toward the actin plus-end. The magnitude of the sum of the vectors for all actin filaments that could potentially contact the liposome was defined as the LPA (Fig. 5A) (see Methods). Mathematically, this resembles the techniques used to calculate the overall magnetic field polarity resulting from the net effect of many individual dipoles within a metal (Zahn, 1979). For example, if actin filaments in a local network are “not aligned,” the unit vectors cancel when summed, resulting in a net vector with near zero magnitude (Fig. 5A). As actin filaments become more aligned, the net vector magnitude increases, eventually reaching a maximum magnitude when the filaments are aligned, and the plus-ends of the filaments point in the same direction. Normalizing this net vector magnitude, by dividing by the number of actin filaments in contact with the liposome, gave LPA values ranging from 0 to 1 (Fig. 5A).

**Figure 5.**
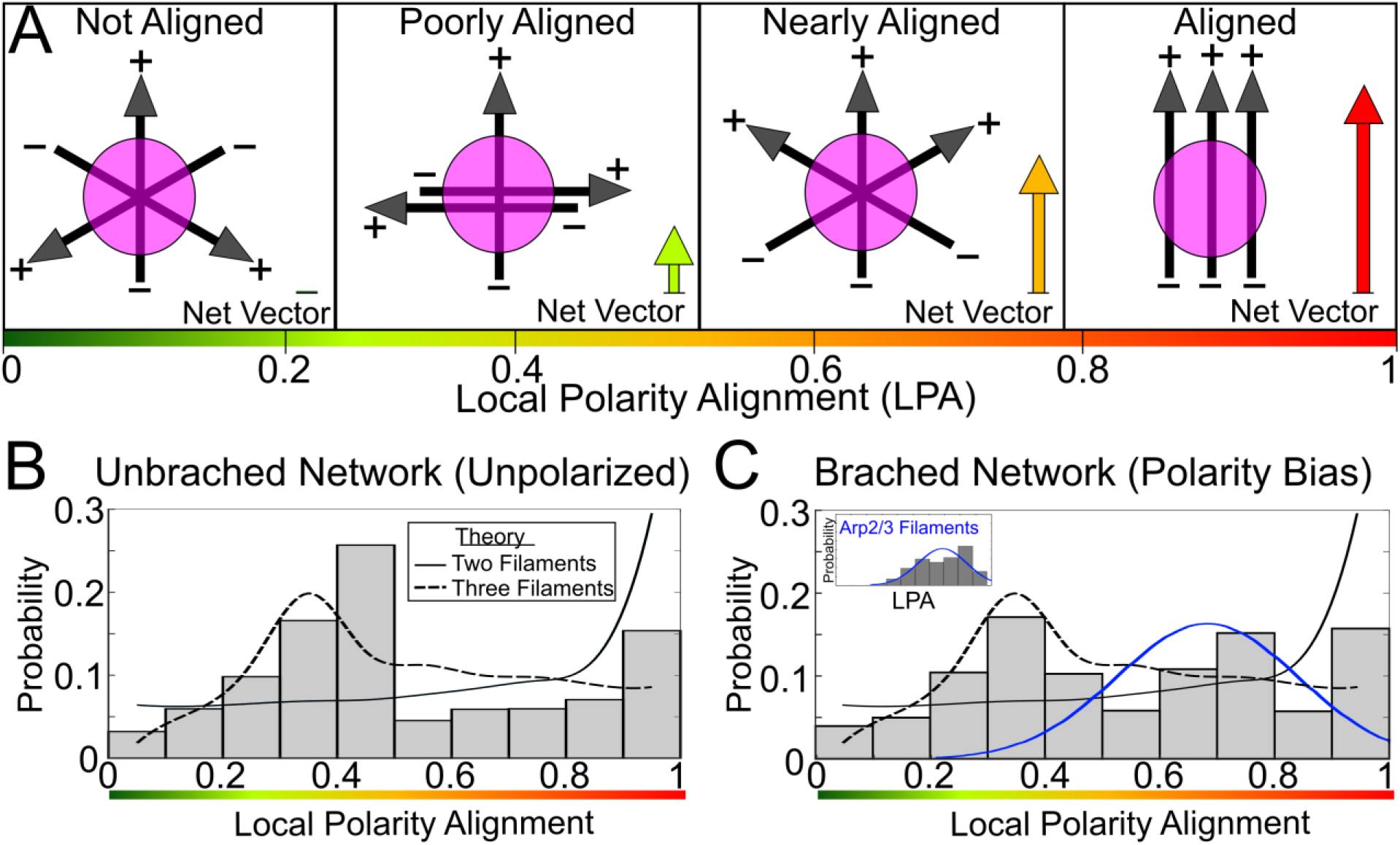
Local Polarity Alignment. **(A)** Schematic of Local Polarity Alignment (LPA) Characterization for Three Actin Filaments. LPA ranges from 0 to 1 (green to red) and is a normalized measure of the actin filaments’ (black arrows) polarity alignment. These filaments are within contact range of the MyoVa motors on the liposome surface (magenta) (i.e. the Local Filament Density; see Methods). Four examples of LPAs are provided. At an LPA of zero, the actin filaments and their plus-ends are at 120 degrees relative to each other. If each filament is treated as a unit vector, their unit vectors sum to a net vector of magnitude zero (i.e. not aligned). As the plus-ends of the filaments become more aligned (i.e. generally point the same direction), the net vector magnitude increases. When all the filaments point in exactly the same direction, the filaments are aligned and the net vector (red) reaches its maximum value. Mathematically, LPA is the magnitude of the net vector normalized by the Local Filament Density. **(B)** Probability distribution for LPA in an unbranched network shows two major peaks, which align with theoretical predictions of two (solid), and three (dashed) randomly oriented filaments (see Methods). **(C)** Probability distribution for LPA for Arp2/3-branched filament networks show the same two peaks seen in (B) with a new third peak between 0.6-0.8, indicating a slight polarity bias has been introduced into the network, as suggested by the theoretical LPA distribution for a local network created solely from Arp2/3-branched filaments (inset, blue line) (see Methods).

The LPA distribution calculated for the unbranched filament networks studied, demonstrated two peaks (Fig. 5B). Since the LPA is related to the LFD (i.e. the number of filaments in contact with the liposome), which was most often between 2-3 filaments (Fig. 3D), we hypothesized that these peaks reflected local contact networks of 2 and 3 filaments, whose actin filament polarities are randomly aligned. Therefore, we calculated theoretical LPA distributions for randomly aligned 2-filament (Fig. 5B, solid curve) and 3-filament (Fig. 5B, dashed curve) local contact networks (see Methods) that show peaks at approximately the observed values. The agreement between the theoretical and experimental LPA distributions for the unbranched networks suggests that randomly aligned 3-filament local contact networks underlie the central peak near an LPA of 0.4, while randomly aligned 2-filament local contact networks underlie the rise in the LPA distribution near an LPA of 1.

In comparison to the unbranched networks, Arp2/3-branched networks showed a third peak (~15% of population) centered at an LPA between 0.6 and 0.8, or ‘nearly aligned’ (compare Fig. 5A and 5C). We hypothesized that this peak arose from local contact networks of increased polarity alignment due to Arp2/3-branched actin filaments. To test this, we created a Monte Carlo simulation to estimate the LPA distribution that would arise from a random network of filaments comprised solely of Arp2/3-branched filaments with their characteristic 70±14° branch points (inset Fig. 5C) (see Methods) (Mullins, Heuser, & Pollard, 1998). Interestingly, this theoretical Arp2/3-branched network is described by an LPA distribution that closely matches the additional peak in the experimentally observed Arp2/3-branched network LPA distribution (Fig. 5C, solid blue curve). Thus, liposomes moving within both the unbranched and Apr2/3-branched networks appeared to encounter local networks of actin polarity alignment ranging the full spectrum from not aligned to fully aligned (Figs. 5B, C). However, liposomes transported within the Arp2/3-branched network were exposed to local contact networks with greater local polarity alignment (Figs. 5B, C), which may contribute to the observed increase in directed and decrease in stationary modes of motion in these Arp2/3-branched networks.

### In Silico Mechanistic model of 3D network transport

To explain the results of our *in vitro* experiments, we used a previously published *in silico* model that simulates the complex interactions between MyoVa motors on the cargo surface and the multiple actin filaments with which they engage (Lombardo et al. 2017). In the model, each 350 nm liposome has 10 individual MyoVa motors that diffuse freely across the liposome surface. Each motor can bind to, take 36nm (or occasional 31nm) steps along, and detach from available actin filaments according to force dependent rates described previously (Clemen, Vilfan, Jaud, et al. 2005; Kad, Trybus, and Warshaw 2008; Lu, Kennedy, Warshaw et al. 2010; Oguchi, Mikhailenko, Ohki, et al. 2010; Veigel, Schmitz, Wang, et al. 2005). Motors are modeled as rigid rods, which diffuse rapidly across the cargo surface to which they are attached via elastic linkages with a torsional stiffness of 0.25 pN nm rad^−1^ and an extensional stiffness of 1 pN nm^−1^ (Lombardo et al. 2017). The action of the MyoVa motors is simulated with a version of the Gillespie algorithm (Gillespie 1977), which accounts for stochastic fluctuations in systems with small numbers of molecules, modified to account for the force-dependence of the motor’s reaction rates. The forces on each actin-bound motor, and the position of the liposome, are determined by finding the mechanical equilibrium between the cargo and motors (Lombardo et al. 2017).

### Local actin position and polarity dictate the mode of liposome motion

The model has no free parameters with which to fit the data; it is therefore predictive. Simulated liposomes were challenged to navigate virtual 3D actin networks with filament positions and polarities identical to those experimentally defined in our microfluidic chambers (Figs. 2, S5, S6). When simulations were performed on networks from the unbranched and the Arp2/3-branched experimental conditions, the model recapitulated the finding that directed transport is favored on the Arp2/3-branched networks (Fig. 3C). Given that the model reproduced our experimental observations, we used it to determine the mechanistic principles underlying the differing modes of motion we observed on the Arp2/3-branched and unbranched networks.

We hypothesized that the 3D position and polarity alignments of actin filaments near the liposome, on the scale of ~1 μm (3-4 liposome diameters), dictate the mode of liposome motion. Specifically, local networks with polarity aligned filaments (Fig. 5A) would promote directed motion, whereas local networks with filaments that are not polarity aligned would promote stationary liposomes. We termed these local actin networks, “actin motifs.” Therefore, Arp2/3-branched networks, with their greater probability of having polarity aligned actin motifs (Fig. 5C), should bias the modes of motion distribution toward directed motion (Fig. 3C). We tested this prediction by replacing a subset (i.e., 37%) of actin motifs in an unbranched network with Arp2/3-branched filaments and in fact, the modeled distribution of alpha values (i.e. modes of motion) for the unbranched network shifted to that of the Arp2/3-branched network (Fig. 6A; see Supplementary Material for simulation details).

**Figure 6.**
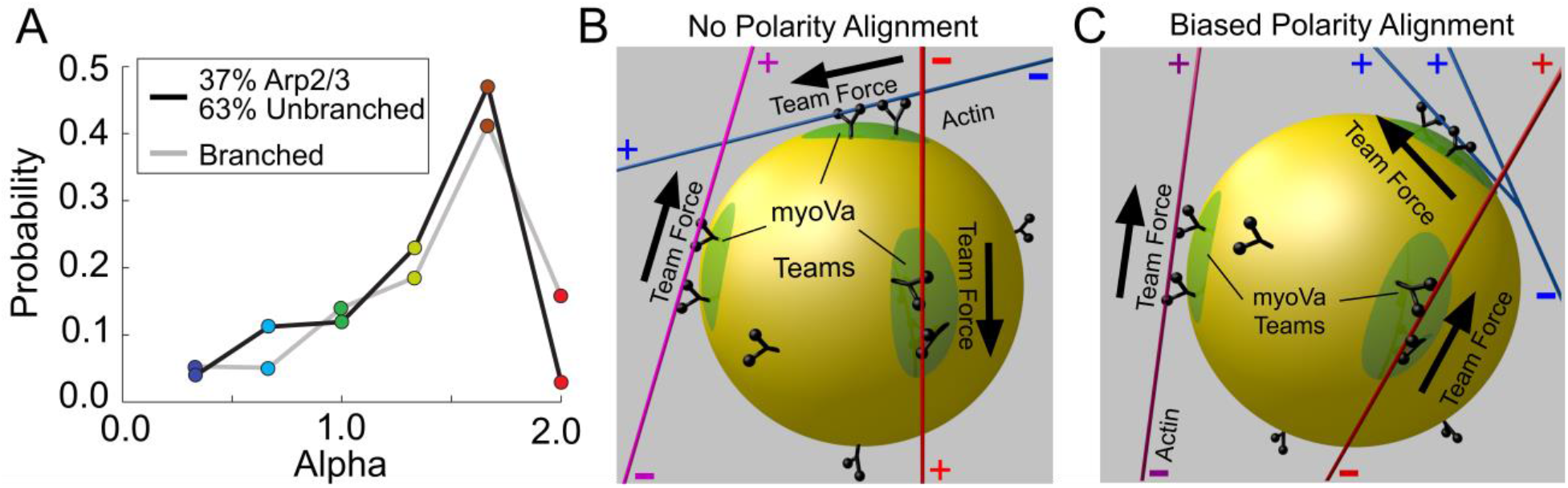
Local Polarity Alignment Determines Net Force Production from MyoVa Motor Teams. **(A)** Alpha histogram for simulations within actin networks where 37% of the unbranched filaments have been replaced with arp2/3 branched actin. Differences between the alpha distributions from simulations shown in Figure 3B&C are mitigated by exchanging a minority (37%) of the actin filaments in the unbranched networks for arp2/3 branches. **(B)** MyoVa motors (black) are recruited to nearby actin filaments (red, blue, magenta) to form teams (green) on the lipid cargo (yellow). Individual motors within the team bind and detach from actin, which creates dynamic teams which can exchange with motors freely diffusing across the cargo’s fluid lipid bilayer surface. When the Local Polarity Alignment is ‘poorly aligned’ or ‘not aligned’, the force (black arrows) produced by each MyoVa team can be directed away from each other producing resistive forces and thus little productive transport. These antagonistic motor teams and their resistive forces can result in stalled MyoVa motors, which then tether cargo to the actin. **(C)** When actin filaments have a biased polarity alignment (nearly aligned or aligned LPA), forces from the different motor teams are generally aligned and directed cargo transport can occur more easily.

To further emphasize the impact of actin motif polarity alignment on the modes of liposome motion, we recreated actin motifs *in silico* from experimental unbranched networks (Fig. S8) in which the experimentally observed trajectories were predominantly either directed (average alpha = 1.51) or stationary/diffusive-like (average alpha = 0.78). We then simulated 50 liposome trajectories on each motif with the liposome starting at the same location within the actin motif as in the experiment. However, the liposome’s azimuthal angle on the actin filament at its starting point are randomized to ensure that the simulated trajectories and their modes of motion are due to the actin motif geometry and not specific to the liposome’s starting azimuthal position. These simulations show that there are significantly more directed trajectories (alpha >1.33; p < 0.001, t-test) on actin motifs where directed motion was observed experimentally (Fig. S9), whereas more stationary/diffusive-like trajectories (alpha<1.33; p < 0.001) occur on actin motifs where stationary/diffusive-like motion was observed experimentally (Fig. S9).

### Motor teams engaged with separate actin filaments generate cooperative or opposing forces

As described above, our *in silico* model results indicate that the modes of liposome motion are dependent on the local actin motif and its filament polarity alignment. This dependence on the actin motif alignment is due to it dictating the directionality of forces generated by individual motors or teams of motors on the liposome surface that are engaged with separate actin filaments within the local network. In fact, the advantage of *in silico* modeling is that the force generated by every actin-engaged motor, whether as a single motor or in a team, is defined and thus, its effective contribution to the resultant mode of motion is defined as well. The most dramatic example being stationary liposomes, which the model shows is the result of a tug-of-war between separate motor teams engaged with two or more filaments with unaligned or poorly aligned polarity. Effectively, each team’s force does not contribute to directed forward motion but rather serves to counterbalance the force of the opposing team during the tug-of-war. This tug-of-war or “pause” in motion will last until the detachment of all of the opposing team’s motors occurs (Figs. S13-S15, Supplementary Movies S8-S10 and SM section 3.3). Interestingly, simulations on actin motifs identical to those observed experimentally in which liposomes paused, resulted in simulated trajectories where liposomes paused within 300nm of the 3D position observed experimentally (Fig S12), further evidence supporting the predictive capacity of this model. Based on this example for the stationary mode of motion, one can then provide a mechanistic basis for directed and diffusive-like modes of motion that are described in more detail below (see Discussion).

## DISCUSSION

Many secretory cells rely on the MyoVa motor’s ability to serve a dual role as both a transporter of, and a tether for, secretory granules as they move through the cytoskeleton to sites of secretion at the plasma membrane (Balasanyan & Arnold, 2014; Rosé et al., 2003; Rudolf et al., 2011; Wu et al., 1998). The goal of this study was to determine how, within a 3D actin filament network, local actin filament position and polarity influence the transport of lipid-bound cargo by teams of MyoVa. Therefore, we created two different 3D actin filament networks, one without (Fig. 5B) and, by introducing Arp2/3-branched filaments, one with (Fig. 5C) inherent local alignment of individual filament polarities. This latter network was designed to mimic cellular networks where MyoVa is known to operate, such as those of the dendritic spine in which the Arp2/3 complex creates a bias in filament polarity toward the cell exterior (Pollard & Cooper, 2009; Wagner et al., 2011). Our study shows that, in random networks created from unbranched actin filaments, the modes of MyoVa-driven liposome transport are approximately equally distributed between stationary, diffusive-like, and directed motions (Fig 3B). However, networks with higher Local Polarity Alignment (LPA, due to Arp2/3-branched filaments) produce more frequent directed transport, even with no change in local actin filament density. This finding suggests that the degree of a network’s actin filament polarity alignment may be matched to the physiological demands for MyoVa-driven cargo transport in specific intracellular domains.

### The molecular basis for modes of cargo motion: Directed motion within a 3D network

Ascribing a molecular basis for stationary, diffusive-like, or directed motion of MyoVa-bound cargo is difficult within the crowded cytoplasmic milieu of living cells, where the presence of dynamic cytoskeletons and many different motor types can all exert separate influences on intracellular cargos. However, in our simplified *in vitro* system, directed motion, averaging microns in length, can only arise from the active stepping of the cargo-bound MyoVa motors along stable actin filament networks. When MyoVa motors on the surface of a liposome encounter multiple actin filaments within their local actin network, these motors have the opportunity to engage the actin, and form teams on each available filament (Figs. 6B,C). If the polarities of the local actin filaments are aligned, the forces produced by these teams of MyoVa motors will also be aligned, leading to cooperative force production (Fig 6C; Movie S2). These forces do not need to be perfectly aligned to maintain directed motion, and we observed long, directed trajectories along Arp2/3-branched filament networks where the LPA was consistently ‘nearly aligned’ (Fig. 7A) (Movie S3). One explanation for this is that the application of slight off-axis forces up to ~45° (relative to the direction of MyoVa motion toward the plus-end of actin) have been shown to have little influence on MyoVa motor’s processivity or stepping kinetics (Oguchi et al., 2010). Thus, MyoVa motors are not sensitive to the slightly off-axis loads produced by the neighboring motor teams on the liposome surface. This may explain why MyoVa motors transport their cargo long distances along Arp2/3-branched networks to the tip of dendritic spines in Purkinje neurons (Wagner et al., 2011) and with straight trajectories in the keratinocyte lamellipodia (Hariadi et al., 2014).

**Figure 7.**
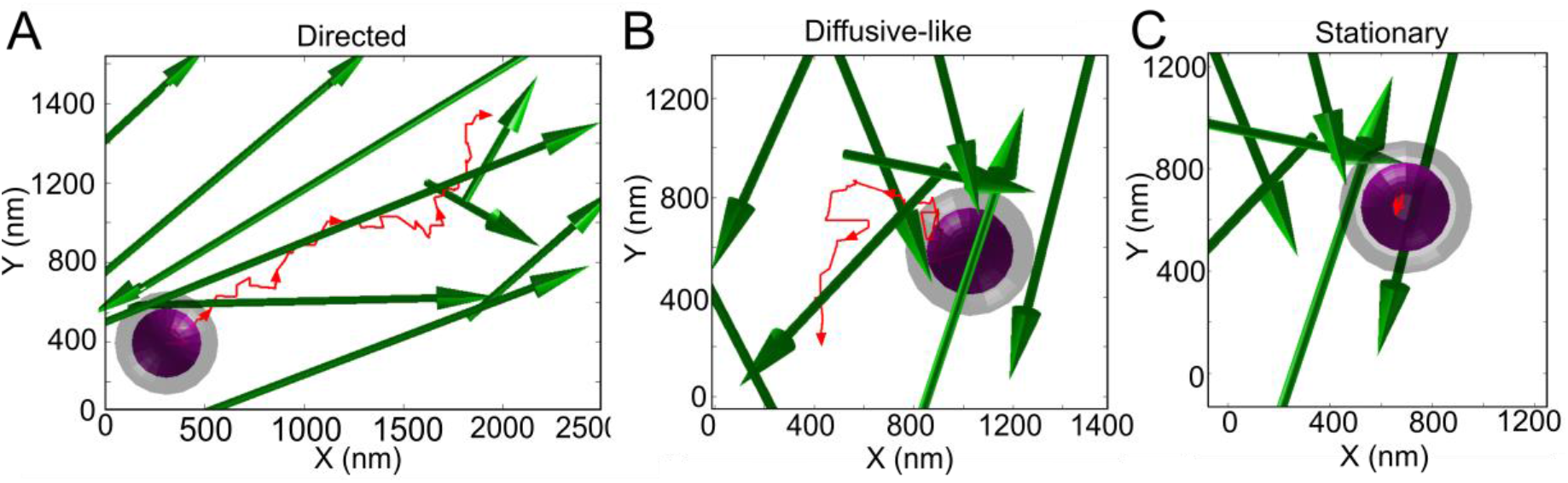
Actin Filament Polarity Alignment Affects Liposome Transport. **(A)** Representative directed trajectory (red) from a 350 nm diameter liposome (purple) within an Arp2/3-branched network with actin filaments shown (green; plus-end shown by arrow head). Grey ‘halo’ sphere around liposome represents the theoretical ‘reach’ of the surface-bound MyoVa (see Methods). The LPA (average=0.8) for the actin filaments at every time point is defined as ‘nearly aligned’. Actin was imaged using 3D STORM and plotted using techniques shown in Figs. 2 and 4 (Movie S3). **(B)** Representative diffusive-like trajectory from an unbranched network. The liposome demonstrates diffusive-like motion as its MyoVa meander along an actin network. Most of the liposome’s time is spent on actin filaments in the middle range of LPA (Average: 0.59) between ‘poorly aligned’ or ‘nearly aligned’ (Movie S4). **(C)** Representative stationary trajectory from an unbranched network. When challenged with multiple filaments whose polarities point in opposing directions (LPA: 0.2), the liposome is stationary as result of an unresolved tug-of-war between competing motor teams that resist each other (Movie S5).

### Diffusive-like motion within a 3D network

In cells, when cargos undergo diffusive-like modes of motion, true diffusion presumably occurs when the liposome is disengaged from the cytoskeletal network. However, a cellular cargo can also meander processively by constantly switching filaments in areas of poor filament polarity alignment (Nelson et al., 2009; Pierobon et al., 2009). The observed movements of the diffusive-like liposomes in our unbranched networks are 24x slower (diffusion constant = 0.059±0.002μm^2^/s) (Table S1) than predicted for a 350 nm liposome in water (i.e. ~1.4 μm^2^/s). This measured diffusion constant is also lower than would be expected from a cargo without motors that is entangled in an actin network with similar Local Filament Densities (LFDs) we report here (Gardel, Valentine, Crocker, Bausch, & Weitz, 2003). Therefore, the molecular basis for diffusive-like motion observed in our networks is the meandering motion produced by teams of motors within local regions of poorly aligned polarity (Figs. 6B, 7B, Movie S4). This conclusion is supported by our simulations, where the tracking of individual motors in diffusive-like trajectories shows mostly processive stepping interrupted only by quickly resolved (<1s lifetime) tug-of-wars (Movie S5). In the cell, this diffusive-like motion may not be futile. MyoVa-driven meandering of cargo along actin networks following long range transport by kinesins and dyneins may provide a dispersive mechanism for cargo to sample the cell membrane domain for target binding sites (Gross et al., 2002; Kögel et al., 2010). Studies of pigment granules in Xenopus melanophores support this concept where the distance traveled by a melanosome correlated negatively with the number of times the cargo switched tracks, reducing directed motion to apparent diffusive-like wandering (Gross et al., 2002).

### Stationary mode of motion within a 3D network

The presence of stationary trajectories in both unbranched and Arp2/3-branched actin filament networks (Fig. 3B,C) could arise from two possible molecular mechanisms: 1) A tug-of-war arising from multiple force producing motors bound to separate filaments which are not polarity aligned, as described above (Figs. 6B, 7C) (Movie S6 & Movie S7); 2) Liposome entanglement, where the filaments themselves serve to cage the liposome, independent of motors attaching to the actin filaments. In cells, these two mechanisms are not mutually exclusive and may be complimentary toward creating stationary liposomes within the networks. Entanglement within an actin filament network would be dependent on the local actin filament density, with denser networks promoting more stationary modes of motion (Gardel et al., 2003). However, the local filament density (LFD) in the two networks studied here was nearly identical (Fig. 3D), suggesting that the greater frequency of stationary liposomes in the unbranched, random polarity aligned networks is due to the tug-of-war between motor teams, as suggested above and supported by the tracking of motors in our simulations (Movie S7).

When motors engage in a tug-of-war, they apply forces on each other (Fig. 6B) (Hendricks et al., 2010, Ali et al., 2011, Gao et al., 2018). An understanding of how a tug-of-war between teams of motors can result in stationary liposomes can be derived from single MyoVa studies in which opposing loads are applied. Specifically, with resistive loads, the motor takes slower and less frequent forward steps and more frequent back steps until the motor stalls when the resistive load equals the motor’s maximum force generating capacity (Clemen et al., 2005; De La Cruz et al., 1999; Kad et al., 2008). Consequently, resistive forces between motor teams interacting with actin filaments in a poorly polarity aligned network can induce tethering of cargo within the network (Fig. 6B, 7C) (Movie S6, Movie S7). Indeed, when simulated liposomes being transported on one actin filament by a MyoVa team encounter a second actin filament, they engage in longer tug of wars, and therefore exhibit longer pauses, the less the two actin filaments are aligned (Fig. S16). Similar results have been observed experimentally by kinesin driven cargos at microtubule intersections (Bergman et al., 2018). Such MyoVa-induced tethering might explain the ‘reserve pool’ of stationary insulin granules within pancreatic beta cells that has been observed ~400-600nm from the plasma membrane within disorganized actin networks (Heaslip et al. 2014; Rorsman & Renstrom, 2003).

### Conclusions

Within the cell, there exists a vast cellular toolbox of actin binding proteins that can manipulate the 3D position and polarity alignment of actin filaments within the cytoskeletal network. The dynamic and malleable nature of 3D cellular actin structures and polarities provide a means for the cell to quickly manipulate the production of opposing vs. cooperative forces from separate MyoVa motors or teams of motors on the same cargo. Additionally, since motors only detect the polarity alignment of the filaments with which they are in contact, the cell can then organize actin filament networks to achieve spatial control for specific cellular transport processes (Fig. 1A). This presents a potential mechanism where the various modes of vesicle motion can be spatially fine-tuned in time and space (Heaslip et al., 2014; Hendricks et al., 2010; Kapitein et al., 2013; Langford, 2002) (Fig. 1A). Future studies will need to explore more dynamic actin filament networks, where actin networks can be deconstructed through actin severing proteins and rebuilt through polymerization-enhancing proteins so that filament position and polarity alignment can modulate a cargo’s modes of motion in real time. Additionally, it is conceivable that the principles presented here are not limited to MyoVa motors and are utilized broadly in both actin- and microtubule-based transport systems as a robust regulatory mechanism to control transport. The incorporation of mixed networks with both microtubules and actin filaments with their respective motors bound to the same cargo will provide insight into the interplay between long range transport by microtubule-based motors and final delivery by myosins.

## METHODS

### Liposome preparation, characterization, and motor attachment

Liposomes were prepared as described extensively in Lombardo et al. (2017), which has been summarized for clarity and repeatability below. We used extrusion through a polycarbonate 25mm membrane (Thermo-Fisher #097322) to create 350 nm diameter phospholipid liposomes, composed of (molar ratio) 84 parts DOPC (1,2-dioleoyl-snglycero-3-phosphocholine), 5 parts PEG-ylated phospholipid (1,2-dioleoyl-sn-glycero-3-phosphoethanolamine-N-[methoxy (polyethylene glycol)-2000], 5 parts cholesterol, 5 parts 1,2-dioleoyl-sn-glycero-3-phosphoethanolamine-N-[4-(p-maleimidophenyl)butyramide] (MBP:PE) and 1 part carbocyanine dye DiI or DiO Cell-labelling Solution (Thermo Fisher Scientific) (Nelson et al., 2014). Liposomes were incubated with a final concentration of 1mM thiolated-Neutravidin (SH-NaV) and incubated at room temperature for 1 hour to covalently conjugate the maleimide moiety of the MBP:PE within the liposome membrane to the thiol groups of the SH-NAV (Nelson et al., 2014). Centrifugation at 400,000xg for 10 minutes was used to remove excess SH-NAV. The liposomes were then resuspended in Phosphate Buffered Saline (PBS) (137mM NaCl, 2.7mM KCl, 10mM Na_2_HPO_4_, 1.8mM KH_2_PO_4_, pH 7.2) and extruded through filter membranes of pore size 650 nm (Thermo-Fisher #097322), which resulted in a final liposome diameter of 350.1±31.9 nm (mean±s.d) when measured by dynamic light scattering using a Wyatt Technology DynaPro model MSX-TX and Dynamics V6 software (Lombardo et al., 2017).

A double-headed, heavy meromyosin myosin Va (MyoVa) construct was expressed with calcium-insensitive calmodulin light chain using a baculovirus/Sf9 cell system and incorporated with an 88-aa biotin ligase recognition sequence at the C-terminal end (Hermanson, 2008). The construct was purified using affinity chromatography to a C-terminal FLAG tag on the myosin heavy chain (Hodges et al., 2007; Krementsov et al., 2004). The C-terminal biotin on the MyoVa construct bound to the SH-NaV on the liposomes, which allowed us to attach the MyoVa motors to the liposome exterior surface (Lombardo et al., 2017). To do this, MyoVa motors (3.3μl, 500 nM) were diluted into 6.7 μl of actin buffer (AB) (25 mM imidazole, 4 mM MgCl2, 1 mM EGTA, 25 mM KCl, 10 mM DTT, with 3.5 mg·ml^−1^ glucose, 40mg·ml^−1^ glucose oxidase, 27 mg·ml^−1^ catalase, 100mg·ml^−1^ creatine phosphokinase, 1 mM creatine phosphate) with 1 mg·ml^−1^ bovine serum albumin (BSA) and then conjugated to 350 nm DOPC liposomes by mixing 10 μl of 3.9 nM of the liposomes into the diluted MyoVa and incubating for 15 min. at room temperature.

To measure the number of MyoVa motors conjugated to each liposome, we used a MyoVa construct that was identical to the MyoVa construct described above except that it included an N-terminal Yellow Fluorescent Protein (YFP) on each motor domain (YFP-MyoVa). We employed a fluorescence photobleaching approach, originally developed by Nayak and Rutenberg (Nayak & Rutenberg, 2011), and one we applied previously to an identical liposome system (Lombardo et al., 2017). Briefly, the intensity decay of the fluorescent YFP-MyoVa, bound to the liposome surface, were recorded in Total Internal Reflection Fluorescence (TIRF) while liposomes were in contact with the glass surface. The integrated fluorescence intensity of each liposome was measured as the intensity decayed over time as the YFP fluorophores on the YFP-MyoVa photobleached. The deviation on the rate of fluorescent decay was then related to the intensity of a single YFP under the conditions imaged using the technique described in Nelson et al. (2014). Knowing the intensity of the individual YFP fluorophore allowed us to calculate the total number of fluorophores and thus, YFP-MyoVa bound to the liposome from the initial fluorescent intensity before any photobleaching occurred as shown in Lombardo et al. (2017).

### 3D actin filament network motility assay

Chicken skeletal actin was used for all experiments and prepared as described previously (Pardee & Spudich, 1982), with additional steps for the Arp2/3-branched actin as follows. Fully polymerized filamentous (F)-actin was dialyzed with agitation against G-buffer (2.0 mM Tris Base, 0.2 mM CaCl2, 0.005% NaN3, 0.2 mM Na2ATP, 1.0 mM DTT) for three days, with buffer changes every 24 hours to produce monomeric globular (G)-actin. Once produced, G-actin was clarified by certification at 400,000xg for 20 min and stored at 4°C for no more than 1 week or at −80°C for up to 1 year. Branched actin filaments were produced on the day of experiments by mixing to a final concentration of 5 μM G-actin in a total volume of 30 μl of polymerization buffer (10 mM Imidizole, 150 mM KCl, 3.0 mM MgCl2, 10 mM DTT, 1 mM ATP, pH7.5) at room temperature for 6 minutes. After the initial polymerization, nucleation of branched filaments was initiated by bringing the total volume to 80μl of polymerization buffer with final concentrations of 5.8 μM VCA domain of Westcott Aldrich Syndrome Protein (Cytoskeleton, Inc.), 150 nM of Arp2/3 (Cytoskeleton, Inc.), 2.5 μM of Alexa-647 labeled phalloidin (Thermo-Fisher), 1.6 mM NaATP, and additional G-actin to maintain the total actin concentration of 5 μM and provide monomers to elongate nucleated branches. The mixture was allowed to incubate at room temperature for an additional 30 minutes before being stored on ice. The resultant solution was then diluted with (AB) to a final concentration of 3 μM of F-actin.

Actin filaments were suspended in 3D between silica beads as described previously (Lombardo et al., 2017) with the following changes: 1) Silica beads of varying diameter (500 − 3000 nm in 500 nm increments) were used instead of only a single diameter; 2) Actin was introduced into the flow chamber at a higher concentration of 3 μM to create denser filament networks. To briefly summarize, the sequential steps for creating the actin filament network within the flow chamber as follows. The mixture of multi-sized silica beads was created by mixing a ratio 1:1 for all beads of diameter between 1500 nm and 3000 nm. Beads of 500 nm, 1000 nm were mixed at a ratio of 0.33:1 and 0.5:1 respectively into 1 M TRIS pH 8.0 buffer. The mixture was then incubated with poly-L-lysine as described in Lombardo et al. (2017) and then introduced into a customized 30 μL flow chamber at an approximate concertation of 0.01% solids. This chamber was flushed with AB with 1 mg·ml^−1^ BSA and allowed to incubate for 2 minutes. Then, either the Alexa-647 phalloidin-labelled unbranched, or Arp2/3-branched actin filaments (3 μM) in AB were flowed into the chamber. The actin was incubated for 2 minutes before gently being washed out using AB. The final wash was completed using a specific AB-STORM buffer, which was AB buffer with 77 mg·ml^−1^ of beta-mercaptoethylamine and 1 mM NaATP, which was used for all STORM imaging (AB-STORM buffer). The imaging of liposomes was done in AB and 1 mM NaATP. This buffer was identical to the AB-STORM buffer without the mercaptoethylamine. To enhance the fluorescent imaging of the MyoVa-coated DiO-labelled 350nm liposomes, we first diluted them 400x to a final concentration of 50 pM before introduction into the flow chamber, to encourage sparse attachment to the actin networks. After imaging the 350nm liposomes, a 3x volume wash of AB buffer with 1 mM NaATP was used to remove the 350nm liposomes with the fluid exchanged as gently as possible. To determine the polarity of the filaments (see, Polarity Reporter Cargo Preparation and Filament Polarity Identification) DiO-labelled reporter cargo (i.e. liposomes) were introduced into the flow chamber at a final concentration of 1 nM of MyoVa motors, and imaged as they walked along single filaments. As a control to ensure that the flow of buffer into and out of the chamber had not disturbed the actin, we confirm the stability of actin filaments within the networks. To do this an additional 3x wash of AB buffer was used to remove the reporter cargo, which was followed by the AB-STORM imaging buffer, and then an additional series of STORM imaging on the same field of view (Fig. S2).

### Microscopy, 3D image acquisition and calibration

3D STORM images were acquired using a Nikon N-STORM super-resolution microscope system with excitation of Alexa-647 phalloidin-labelled actin by 647 nm and 405 nm lasers. A cylindrical lens was added to the light path to introduce an astigmatism and allow for 3D imaging as described previously (Huang et al., 2008; Lombardo et al., 2017). Approximately 30,000 images were collected to generate the actin super-resolution 3D reconstruction. Raw TIFF images were then imported into the ImageJ plugin Thunderstorm (Ovesný, Křížek, Borkovec, Švindrych, & Hagen, 2014). Minimum and maximum intensity thresholds were determined on a chamber-to-chamber basis, along with background and localization uncertainty measurements. Fluorophores with an axial ratio greater than 1.3 were eliminated. A 532 nm laser was used to excite fluorescent DiO-labelled liposomes navigating the actin filament networks. The Nikon ‘perfect focus’ system was applied to ensure Z-dimension imaging stability and alignment of the liposomes and actin within the same Z-plane over the duration of any individual imaging session. Thunderstorm cross-correlation drift correction was applied to account for XY drift. Additionally, the silica bead supports were visible as fiducial markers in all imaging channels and were used to cross-correlate between the actin images, 350 nm liposomes, and polarity reporter cargo. Using the ImageJ plugin PIV (Tseng et al., 2012), the fiducial markers could be aligned with sub-pixel resolution. Collectively, these techniques allowed for precise sub-pixel tracking of drift in X, Y and Z both within, and between all actin, liposome, and polarity reporter imaging. To correct for chromatic aberrations, a Z-position lookup table was created using multi-color beads adhered to the glass surface. The lookup table was generated by stepping beads through different Z-positions (±400 nm) using the piezo stage, while imaging them in different color channels. To correct for aberrations arising from differences between the index of refraction between the glass surface and buffer, a rescaling factor was applied to Z-positions as described previously (Lombardo et al., 2017).

### 3D liposome position

The liposome 3D position calibrations shown and described extensively in Lombardo et al. (2017) were used in this study. In brief, raw TIFF images were imported into ImageJ and a two-axis elliptical Gaussian was used to fit the images of fluorescent liposomes. To create a Z-calibration curve, actin was attached to the glass surface of a flow chamber coated with Poly-L-Lysine. 350 nm liposomes identical to those used in all experiments were then flowed into the chamber in AB buffer with 0mM NaATP, causing the liposome-bound MyoVa to attach to the actin without letting go (i.e. rigor conditions for the actin-bound myosins). The fluorescent liposomes on the glass surface were then stepped from −400 nm to +400 nm using a piezo stage. Vertical and horizontal axis fits to the fluorescent liposome images were normalized and then matched to the Z-calibration curve as described previously (Henriques et al., 2010). Precision of the liposome localization was measured by sparsely binding 350 nm liposomes to single actin filaments suspended between silica beads off the glass surface in AB buffer with 0 mM NaATP. This condition represents the uncertainty in localization precision, which results from the combined movements of the suspended filaments itself, and the liposome’s attachment through the myosin motors. The resultant localization precession for suspended 350nm myosin-bound liposomes was 17 nm in X, 18 nm in Y and 30 nm in Z.

### Polarity reporter cargo preparation and filament polarity identification

Smaller, 50 nm, liposomes were produced from the 350 nm liposomes described above and were used as reporters for actin filament plus-end polarity. To do this, 500 μl of 350 nm liposome were sonicated using a 550 Sonic Dismembranator (Thermo-Fisher) for 5 minutes with 0.5 second bursts while kept at 4° C by submersion in an ice bath. The 7x decrease in diameter leads to an approximate 50x increase in solvent exposed surface area of the liposomes. Therefore, incubation with myosin under identical conditions as the 350nm liposomes (3.3 μl of 500 nM motors, 10 μl reporter liposomes, and 6.7 μl AB buffer) lead to a theoretical average of 1 myosin bound for every 5 reporter liposomes. These limiting motor conditions were confirmed through titration of the MyoVa motors, where reduction in the concentration of incubated motors less than 1 motor: 5 liposomes, eliminated observed transport of the reporter cargo along actin filaments.

Reporter cargo were introduced into the actin networks following 350 nm liposome imaging (Fig. 4). This prevented the small chance that a photobleached, defective, or unseen reporter cargo could influence the trajectories of the 350 nm liposomes within the actin networks. Trajectories of the reporter cargo were aligned to the actin STORM images using the silica bead supports as fiducial makers and employing the same ImageJ-based PIV cross-correlation analysis used for the 350nm liposomes. Once aligned in X and Y, the raw TIFF images of the reporter cargo’s fluorescent signal were overlaid onto the STORM reconstruction of the actin filaments to make a movie of the reporter cargo transport along the network filaments (Fig. 4) (Movie S1). The reporter cargo’s fluorescent signals were then tracked using the sub-pixel localizing ImageJ plugin (MTrackJ). The reporter cargo trajectories were required to be a minimum of 150 ms in duration to be considered the result of single MyoVa undergoing processive movement. The direction of travel was then considered to be towards the plus-end of the actin filament. In this way the polarity of the actin filament was identified. Filament polarity analysis was performed once actin filaments were localized within the network following STORM reconstruction (see below). By this approach, virtually every actin filament’s polarity within the network was identified.

### Actin filament 3D position fitting

Each individual Alexa-647 3D STORM localization (see, Microscopy, 3D Image Acquisition and Calibration) was saved as an entry in a .csv file. Full frame data sets were cropped to regions of interest, of up to 120 filaments, then imported into the statistical programing language R. The regions of interest were plotted on a dynamic 3D plot, which could be rotated freely and observed from any angle using the RGL package library (Fig. 2) (Adler et al., 2014). Initial starting inputs for the filament fitting were made by manually identifying a potential start and end point for each individual filament. The fitting algorithm then created a tubular volume with a 200 nm radius between the start and end points and collected all STORM localizations within the volume (Fig. 2B). A least squares linear regression was then fit through the 3D STORM localizations, creating a best fit line segment for each actin filament. The ends of the line segment were allowed to elongate or shorten to best fit data that continued beyond or ended before the initial estimates. This was limited to a total length change of (400 nm) to prevent fitting to data from other sources (e.g. a silica support bead), which were beyond the end of the fitted filament. Each filament’s fit was then tagged with an ID number and saved as a line segment defined by two points, each with an X, Y, and Z position, where the filaments measured position spanned between the two points. The root mean squared deviation between the fit and the STORM localization data was then used to measure the precision of the actin fit. The polarity of each filament was then added to these fits as a binary identifier indicating which of the two points was the plus-end of the actin (see above).

### Mean squared displacement (MSD) and change point analysis

Mean squared displacement analysis originally described in Heaslip et al. (2014) was adapted to accommodate 3D data for this study using the equation below.

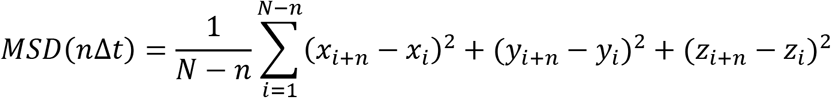

Where N is the total number of frames in the trajectory, n is the number of frames for different time intervals, Δt is the time interval between frames for the trajectory (50 ms for 350 nm liposomes), and (xi, yi, zi) is the position of the liposome at time i. A custom “change point’ algorithm was written in the statistical analysis program R, which segmented trajectories into “directed”, “diffusive-like”, and “stationary” regions. The segmentation was done by calculating a rolling-window MSD, which allows for an unbiased identification of distinct modes of motion within single trajectories. The modes of motion are identified by plotting the MSD versus time (ΔT) on a log-log axis (Fig. 3A inset) and calculating a linear fit through the plotted data. The slope of the fit line, defines the diffusive exponent, alpha value (α). The alpha value can range between 0 and 2 with an α of 0 defining completely stationary motion, α of ~1 would arise from diffusive-like motion and an α of 2 from purely directed motion (Saxton & Jacobson, 1997). We defined, stationary, diffusive and directed motions as three equal ranges of alpha values as follows: stationary (α≤0.67), diffusive (0.66 < α < 1.33), and directed (α ≥ 1.33). These cutoffs were set by analysis of known stationary particles imaged using the same optical system as our experiments, which resulted in α=0.17±0.09 defined previously (Heaslip et al., 2014). To identify transitions between modes of motion we summed the Bayesian Information Criterion (BIC) for MSD fits where:

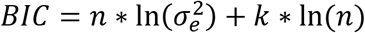

Where k is the number of parameters for the fit (in this case 4), n is the number of measurements in the fit and σ_e_ is the error in the fit to the MSD data described above. As the fit varies from the MSD vs. time data, it indicates a mode of motion changed, and the BIC value drops. BIC transitions between modes of motions, diffusion coefficients, run lengths, and associated error terms were calculated as described in Heaslip et al. 2014. Statistics between alpha values in the Arp2/3-branched and unbranched networks was accomplished using the Kolmogorov-Smirnov test. Stationary lifetimes, diffusion coefficients, directed run lengths and directed velocity measurements were calculated as described in Heaslip et al. (2017). Diffusion coefficients show a log-normal distribution and are thus reported as a mean ± error factor. The error factor is defined as the square root of the width of the 90% confidence interval (Rausand & Høyland, 2004, Heaslip et al., 2017)

### Local filament density and local polarity alignment analysis

We created a theoretical spherical volume, which centered on the liposomes localized 3D position to estimate the number of filaments that MyoVa motors bound to the exterior of a liposome could potentially engage. The theoretical volume was created using constants, which were either experimentally measured in this study or previously published as follows: The diameter of the sphere was the sum total of the 350 nm measured liposome diameter (Lombardo et al., 2017), 100 nm for the approximate length of two (one MyoVa on each side of the liposome) of our MyoVa motor constructs (Trybus, 2008), and the maximum propagated error for the actin localizations and liposome tracking positions (36nm) (Figs. 2A-C, 6) (Lombardo et al., 2017). We centered the volume around the 3D trajectories of the 350 nm liposomes, which had been aligned (see above) to the actin fits on a frame to frame basis (Figs. 6). Any actin localization fits, which passed through this volume were then considered to be in contact with the liposome at that single frame and thus, defined the Local Filament Density for that liposome at that frame. Kolmogorov-Smirnov (K-S test) was used for statistical analysis of the on Local Filament Density between conditions.

The identifier of each contacted filament was stored for each frame and a Local Polarity Alignment (LPA) was calculated from only the filaments, which were considered to be in contact with the liposome. To calculate LPA, we created a spatial vector, which matched the 3D position of each actin filament’s 3D STORM localized fits with a plus-end polarity determined from the reporter liposomes. The magnitude of each vector was set to 1, so that each filament was represented as a unit vector that indicated its polarity. The vectors, for filaments in contact with a 350nm liposome, were then added to each other using simple vector algebra to create a net vector (Fig. 5A). Filaments, which had their plus-ends pointing in opposite directions, would thus subtract from each other and cancel out leading to a net vector of zero magnitude (Fig. 5A). On the other hand, filaments, which pointed in the same direction would add to each other, increasing the net vector’s magnitude. The resulting net vector related information only pertaining to the filaments’ polarity relative to each other, not their physical separation in 3D. Using this analysis, the total magnitude of the net vector would thus be a measure of the alignment of the actin filaments contacting the liposome at any given frame. The LPA was normalized by then dividing the net vector’s magnitude by the number of filaments. To model how the LPA value would be distributed for two filaments over all possible positions and polarities (Figs. 5B,C), we defined two-unit vectors (i.e. actin filaments), each identified by their start (minus-end) and end (plus-end) 3D coordinates. The vector coordinates were first set to be identical. Then, one of the vectors was rotated around its center in the XY plane while the other remained stationary. LPA was then calculated in one-degree increments as it rotated 0 to 360 degrees. The LPA values were then plotted as a histogram and a smoothed line drawn through the histogram. This theoretical distribution was then overlaid onto the data of LPA measurements from the unbranched and Arp2/3-branched filament networks using MATLAB-version R2017A (Figs. 5B,C). This process was similarly repeated for three filaments by rotating all combinations of the three filament vectors and recording the resulting LPA (Figs. 5B,C).

We also modeled the LPA for an Arp2/3-branched filament that had one or two branches in addition to the mother filament that could be contact with the liposome (Fig. 5C, inset). Therefore, a Monte Carlo simulation was created to determine the distribution of LPAs as follows. The simulation assumed that the mother filament was represented as a unit vector. This vector was assigned as the reference point for the addition of new branched filaments within the simulation. The angle at which Arp2/3 branches came off the mother filament was randomly assigned from a normal distribution based off measurements from Mullins et al. (2008), which was reported as 70±14°. We termed this angle the ‘branch angle’. Either one or two filament branches were then created, which were oriented with their plus-end pointing in the direction of the mother filament’s polarity. This was repeated 1000 times for one and two branches, generating 2000 LPA values. These values were then plotted as a histogram, which showed a broad Gaussian distribution centered at a LPA of 0.68 (Fig. 5C, inset). This fit was then plotted on the LPA histogram plot of the experimental Arp2/3-branched filament network (Fig. 5C). The Monte Carlo simulation and plotting were done in MATLAB-version R2017A.

## ACKNOWLEDGEMENTS

This research was funded by NIH T32HL076122 (A.T.L.), Vermont Space Grant Consortium under NASA Cooperative Agreement: NNX10AK67H (A.T.L., D.M.W.), NIH GM094229 (D.M.W.), NIH GM078097 (K.M.T.), NSF DMS1413185 (S.W.). We would like to thank A. Howe and J. Stumpff, for helpful discussions on data presentation and figure design. We acknowledge E. Krementsova, S. Previs, and T. Sledewski for aid in protein production. Nikon 3D STORM imaging was performed at the Microscopy Imaging Center at the University of Vermont with the support of N. Bouffard and D. Taatjes. We would like to acknowledge the contributions of those who helped to create and distribute open source programs used in this work, including, r-project.org, Inkscape.org, Blender.org, MTrackJ, Thunderstorm, and ImageJ.nih.gov.

