## Supplemental_Materials_and_SM_Appendix for "Myosin Va Transport of Liposomes in Three-Dimensional Actin Networks is Modulated by Actin Filament Density, Position, and Polarity"

Supplementary material to accompany:
### Myosin Va liposome transport of liposomes in an *in vitro* three-dimensional actin network is modulated by actin filament density, position, and polarity

Andrew T. Lombardo, Shane R. Nelson, Guy G. Kennedy, Kathleen M. Trybus, Sam Walcott, and David M. Warshaw

#### 1 Supplemental Table

| Network | Stationary Lifetime (Median ± SEM) | Diffusion Coefficient (Mean ± error) | Directed Run Length (Median ± SEM) | Directed Speed (Mean ± SEM) |
|---|---|---|---|---|
| Unbranched | $4.7 \pm 3.0\ s$ | $5.9 \pm 0.02 \cdot 10^4\ nm^2/s$ | $1190 \pm 340\ nm$ | $322 \pm 61\ nm/s$ |
| Arp2/3-branched | $3.0 \pm 0.8\ s$ | $4.1 \pm 0.02 \cdot 10^4\ nm^2/s$ | $1004 \pm 410\ nm$ | $326 \pm 63\ nm/s$ |

Table S1

#### 2 Supplemental Figures

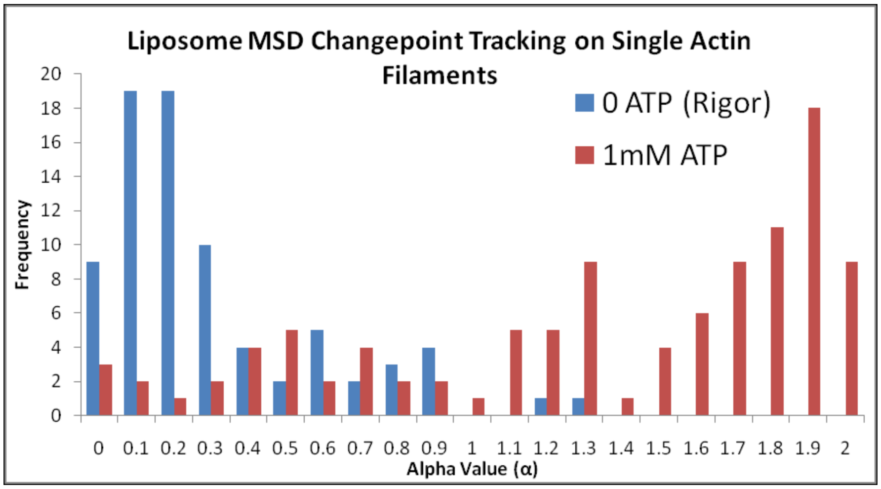


Figure S1: Determination of modes of motion cutoffs. Histogram of liposomes being transported along single actin filaments in rigor conditions (blue) where all liposomes are known to be stationary, and at 1mM (saturating) ATP conditions (red) where robust directed motion has previously been reported (Nelson et al. 2014, Lombardo et al. 2017). Using the alpha value distributions from these conditions, stationary events were defined as $\alpha \leq 0.66$ while directed motion was limited to $\alpha \geq 1.33$. On rare occasions liposomes in the 1mM ATP condition became stationary (red bars $\alpha \leq 0.66$). This observation agrees with previously reported motion of multi-motor-bound liposomes transported on single actin filaments (Nelson et al. 2014).

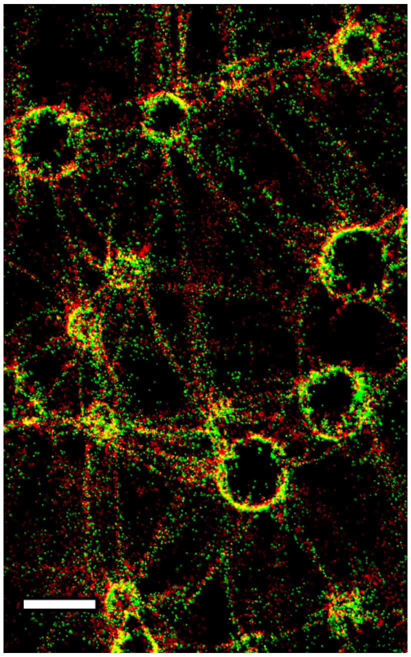

Figure S2: Actin STORM reconstruction overlay before and after liposome and reporter myosin. Actin was imaged using super resolution STORM before introduction of 350 nm liposomes into the flow chamber (Green) as preformed for all other experiments. However, after imaging of 350 nm liposomes navigating the actin and identification of the polarity of the filaments using reporter myosin imaging of the actin under STORM conditions was completed again (Red) identically to the first time. The STORM reconstruction from before and after the liposomes was then overlaid in Imagej to detect if the actin remained stable throughout the entirety of the experiment. Essentially all filaments visible in the first period of actin imaging remained stable and were visualized again in the same location following the experiment, indicating that the liposomes or reporter myosin, or their introduction into the flow chamber did not alter the actin network.

### 3 Supplemental Modeling Section

We have previously developed a 3D model for 350nm-diameter fluid liposomes transported by 10 myosin Va motors (Lombardo et al. 2017). This model quantitatively matches outcome probability (switch, turn, terminate) for fluid liposomes transported through 90 degree actin intersections in 3D. The model also matches the spiral pitch of liposome trajectories, and their average speed along the actin filament. The model has no free parameters with which to fit the data; it is therefore predictive. Here, we will show that (1) the model successfully predicts the differences we observed in the modes of liposome transport motion in unbranched vs. Arp2/3-branched networks, (2) that differences in local actin network geometry underlie these differences, and (3) that force-dependent chemical reactions (i.e., stepping and/or unbinding) of myosin Va are responsible for the model's dependence on local actin network geometry. This text provides additional details, analysis and figures to support the conclusions of the main text. Note: in the main text, we differentiate between the 3D position of actin filaments in the network (measured with STORM) and their polarity (identified with reporter myosin Va). Here, we will use the term "geometry" to refer to both of these quantities.

### 4 Model description

A more detailed description of the model, and further justification of model assumptions, can be found in Lombardo et al. 2017. The system contains three components: the liposome, the actin filaments, and the myosin Va motors on the liposome surface. Each myosin motor undergoes chemical reactions (i.e. stepping forward and backward along actin and detaching from actin) whose rate depends on the force applied to the motor. We determine these forces from a mechanical model of each of the three components.

In the model, actin filaments are rigid rods that are held rigidly in place. Myosin binding sites occur every 5.5 nm along a right-handed double helix, with a periodicity of 72 nm ($\sim$ 14 monomers). Each binding site has a specific orientation, so that a myosin motor, once bound, extends rigidly from the binding site. This preferred orientation, which is orthogonal to the actin filament, rotates azimuthally with the actin periodicity (one full rotation every 72 nm, or 14 monomers).

The liposome is a rigid sphere of radius $R = 175$nm. The surface of the liposome is fluid, so that forces tangent to its surface cause the membrane to flow. Contact between the liposome and the actin filaments is frictionless. There is no attraction or repulsion between the liposome and the actin filaments, other than steric constraints that keep the two from occupying the same space.

Myosin Va molecules are extensible rods, of rest length $\ell = 50$nm, anchored into the fluid surface of the liposome by a deformable pivot. The motor's extensibility is linear, with spring constant $k = 1$pN/nm. The pivot, that connects the myosin motor to the liposome, contains a universal joint with a torsional spring of stiffness $\kappa_\theta = 0.25$pN $\cdot$ nm/rad. There is also a frictionless hinge that allows the motor to freely rotate about an axis normal to the surface of the liposome. The combination of pivot and torsional spring allows us to model the bending stiffness of the myosin molecule. In particular, the spring and pivot work together to cause the motors to extend in the normal direction off of the liposome and resist angular deflections in any direction equally.

Myosin motors can perform forward steps at a rate $k_{step}$, backward steps at a rate $k_{back}$ and detach at a rate $k_{off}$. These rates depend on the component of force on the myosin molecule that points along the axis of the actin filament, $F_x$, which is positive if in the direction opposing forward steps.

The reaction rates are (in $s^{-1}$)

$$k_{step} = \frac{1}{s}\left(\frac{e^{0.04F_x/k_BT}}{13} + \frac{e^{14F_x/k_BT}}{426}\right)^{-1}$$

$$k_{back} = \frac{1}{s} 0.3 e^{1.3F_x/4.14}$$

$$k_{off} = \frac{1}{s}\left(0.1724e^{-1.005F_x} + 0.2634\right)$$

where $k_BT = 4.14\text{pN}\cdot\text{nm}$ is Boltzmann's constant times temperature. All rates come from Kad et al. 2008, but we have introduced a scaling factor, $s$, that allows us to adjust motor velocity to be consistent with our measurements without affecting the motors' run length or stall force. When a motor steps, the position of the myosin molecule is moved forward 36 nm most (78%) of the time, but occasional (22%) 31 nm steps occur. These infrequent short steps cause a single myosin motor to describe a spiral trajectory around the actin filament.

Myosin molecules that are unbound from actin can attach at a rate

$$k_{att} = \frac{k_0}{4\pi R^2}\exp(-\Delta E/k_BT) \tag{1}$$

which follows from the assumptions that diffusion across the surface of the liposome is fast (so the rate is inversely proportional to the surface area of the liposome, $4\pi R^2$) and that attachment occurs via a weak-binding intermediate (so attachment rate depends exponentially on the mechanical energy it takes for myosin to bind, $\Delta E$). To determine this attachment rate, we calculate $\Delta E$ for every binding site on actin that is not already occupied with a myosin molecule. So, for a particular binding site, this requires finding the mechanical equilibrium configuration that would result from a motor binding to that binding site, calculating the energy of that configuration, and subtracting the energy of the current configuration from that value.

Eq. 1 has one free parameter, $k_0$. Rather than specifying $k_0$ directly, it is more convenient to specify an overall attachment rate, which is the sum of $k_{att}$ over every binding site on actin. Generally, this attachment rate depends on the number of myosin motors on the liposome surface that are attached to actin, and the relative position of those attached motors. However, when a single myosin molecule is bound to actin, the liposome is always positioned directly over it. Therefore, we define $k_a = 2.4\text{s}^{-1}$ to be the overall attachment rate when one motor is bound (i.e. the attachment rate of a second motor), which then defines $k_0$ in Eq. 1.

Calculation of the binding energy for the calculation of attachment rates (Eq. 1) is computationally expensive. It requires calculating the equilibrium position of the system for myosin binding to each available actin binding site. Finding the equilibrium position is not trivial, since even assuming mechanical equilibrium, one must simultaneously solve three non-linear equations (force balance in 3D). Further, since the solution is generally not unique, one must identify the correct mechanical equilibrium point. To simplify the calculation, we made the approximation that myosin, as a linear spring, is very stiff. Then, extension of myosin is energetically prohibited. One can then easily find the equilibrium position of the liposome by solving a geometry problem – i.e. if three myosin molecules are bound, where is the liposome such that each myosin molecule is not extended? Solving similar geometry problems for two and one bound myosin, we increase the computational efficiency by roughly $10^3$ – so that simulations that would take overnight can be completed in minutes (see Lombardo et al. 2017 for further justification of this assumption).

To simulate a myosin molecule undergoing these interactions, we use the Gillespie algorithm (Gillespie 1977). Briefly, at each time step, we determine the time of every myosin molecule undergoing every possible chemical reaction (stepping forward, stepping backward, attaching and detaching) by picking random numbers from the appropriate distribution. These times are then sorted, and the reaction with the shortest time is implemented and time is advanced by that time step. In the original method (Gillespie 1977), the next shortest reaction would be implemented, time advanced, and so forth. However, since each reaction generally changes the forces on each attached motor, and since the reaction rate constants depend on these forces, after each reaction occurs we re-calculate all reaction times. Importantly, this method gives the correct stochastic fluctuations that occur when a small number of molecular motors transport a shared cargo.

##### 4.1 Modifications for complex networks

The model, described above, was used to simulate myosin Va motors transporting fluid liposomes through 90-degree, 3D actin intersections (Lombardo et al. 2017). To model more complex actin geometries, we introduced three modifications:

1. Constraints. In the previous model, when in contact with an actin filament, the liposome's movement is constrained, so that it either slides on the filament or it breaks contact. Since the previous experiments only had two actin filaments, we only considered simultaneous contact with zero, one or two actin filaments. Here, we extended the simulations to allow simultaneous contact with up to three filaments. Contact with more than three filaments is prohibited by geometry, given the assumption that both the liposome and the actin filaments are rigid.

2. Actin filament ends. In the previous model, although the actin filaments were finite in length, the liposome never reached or interacted with the ends of the filaments. In these simulations, such interactions are possible. When a motor reaches a filament end, it no longer steps forward, but all other reactions occur as before. When the liposome contacts a filament end, it is constrained to slide on the filament end, or to break contact.

3. Energy check. In these complex networks, geometries often arise where the liposome becomes trapped. That is, even though the myosin Va motors step forward, the liposome cannot follow because its movement is impeded by actin filament(s) acting as a barrier. Our calculation of binding energy only considers the equilibrium position of the liposome, not whether that equilibrium position is accessible. Thus, it is possible that, when the liposome becomes trapped in the simulation, a myosin molecule could attach to an actin filament and the liposome then be placed at an equilibrium position that it cannot achieve without, say, passing through one of the actin filaments. To ensure that such a situation does not arise, we update the position of the liposome by numerically solving the equations of motion for the liposome moving through a viscous fluid, and connected to springy myosin molecules. We then compare the energy of the liposome's final position with the energy predicted by the equilibrium calculation. If the calculated and predicted energies differ by greater than 30%, we do not allow the reaction to occur, and re-run a step of the Gillespie algorithm. Typical differences between calculated and predicted energies are less than 1%.

#### 5 Model predicts experimental observations

We compared the model to our experimental measurements by running simulations on actin filament networks on which we observed liposome transport and starting from initial conditions that matched these experimental measurements. To do so, we had to (1) define the geometry of the actin networks; (2) run multiple simulations from appropriate initial conditions; and (3) analyze the simulations results and compare the results to our observations. We now describe this process.

##### 5.1 Defining the actin network

STORM imaging allows us to accurately measure the 3D position of the actin filaments to great precision, but not all actin filaments are resolved with this technique. As a result, we occasionally see liposome

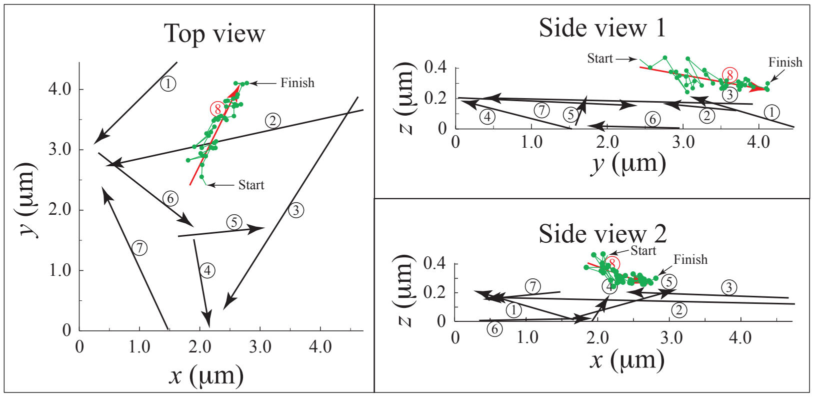


Figure S3: Defining the 3D actin network. Black arrows show the position of actin filaments resolved with STORM, and polarity from the motion of 50nm diameter reporter liposomes. Green shows the trajectory of a liposome, with the start and end of the trajectory labeled, and each dot separated by 0.05s. The position and polarity of an actin filament (red) is inferred by fitting a 3D line to the trajectory.

trajectories that appear directed along an unresolved actin filament. To compare the model to the experimental measurements, we must estimate the position of these unresolved actin filaments. We achieve this by fitting a 3D line through these trajectories (see Fig. S3).

Estimating the position of these unresolved actin filaments is time-intensive, so it was impractical to do it over the entire experimental network. Instead, we focused on two $\sim 4\mu\text{m} \times 4\mu\text{m} \times 0.4\mu\text{m}$ local regions within an *in vitro* experimental network, i.e., one from an unbranched (Fig. S4A) and one from an Arp2/3-branched network (Fig. S4B). In the *in vitro* experiments, we experimentally observed and measured 10 trajectories on each of these local networks. Each local network contained similar numbers of actin filaments (11 for the unbranched, and 12 for the Arp2/3-branched), and similar actin monomer densities (1400 monomers/$\mu\text{m}^3$ for the unbranched, 1900 monomers/$\mu\text{m}^3$ for the Arp2/3-branched).

#### 5.2 Running the simulations

To compare the model to the experimentally observed data, we simulated each of the 10 observed trajectories characterized on each network. For a particular trajectory, we attempted to match the initial condition, starting from a single myosin motor attached, and the liposome oriented as closely as possible to the measured initial condition. We ran the simulation for a longer time than the measured trajectory, and then interpolated to match the simulated times to the measured times. For each experimentally observed trajectory, we ran 10 simulations, since each simulation is different due to the stochastic effects inherent in systems with small numbers of molecules. Note that the actin filaments are discrete and have starting and ending points that represent either where we could resolve them with STORM imaging, or where we observed liposome trajectories. Most of these, we expect, are not true actin filament ends. Therefore, we ensured that none of the simulated myosin Va motors reached and remained at these points.

We analyzed the resulting simulated trajectories using the same changepoint analysis we used for our measurements. Note that if a simulated trajectory was too short to be analyzed by the changepoint analysis program, we discarded that simulation and ran the simulation again. The results for the unbranched network are shown in Fig. S5 and for the Arp2/3-branched network in Fig. S6. In all cases, the agreement between measurement and simulation is reasonable – both in terms of the path taken by the liposome and the alpha value observed along the simulated liposome trajectory.

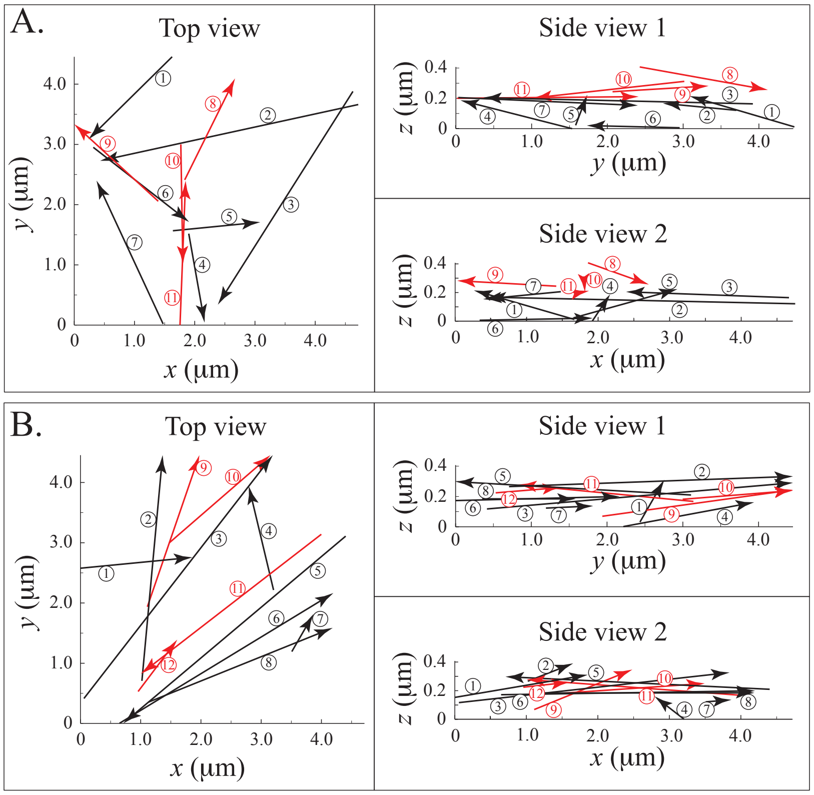


Figure S4: Unbranched (A, top) and Arp2/3-branched (B, bottom) actin networks. Black arrows show the position of actin filaments resolved with STORM, and polarity from the motion of 50nm diameter reporter liposomes. Red arrows show the position and polarity of actin filaments inferred by fitting 3D lines through trajectories (as in Fig. S3).

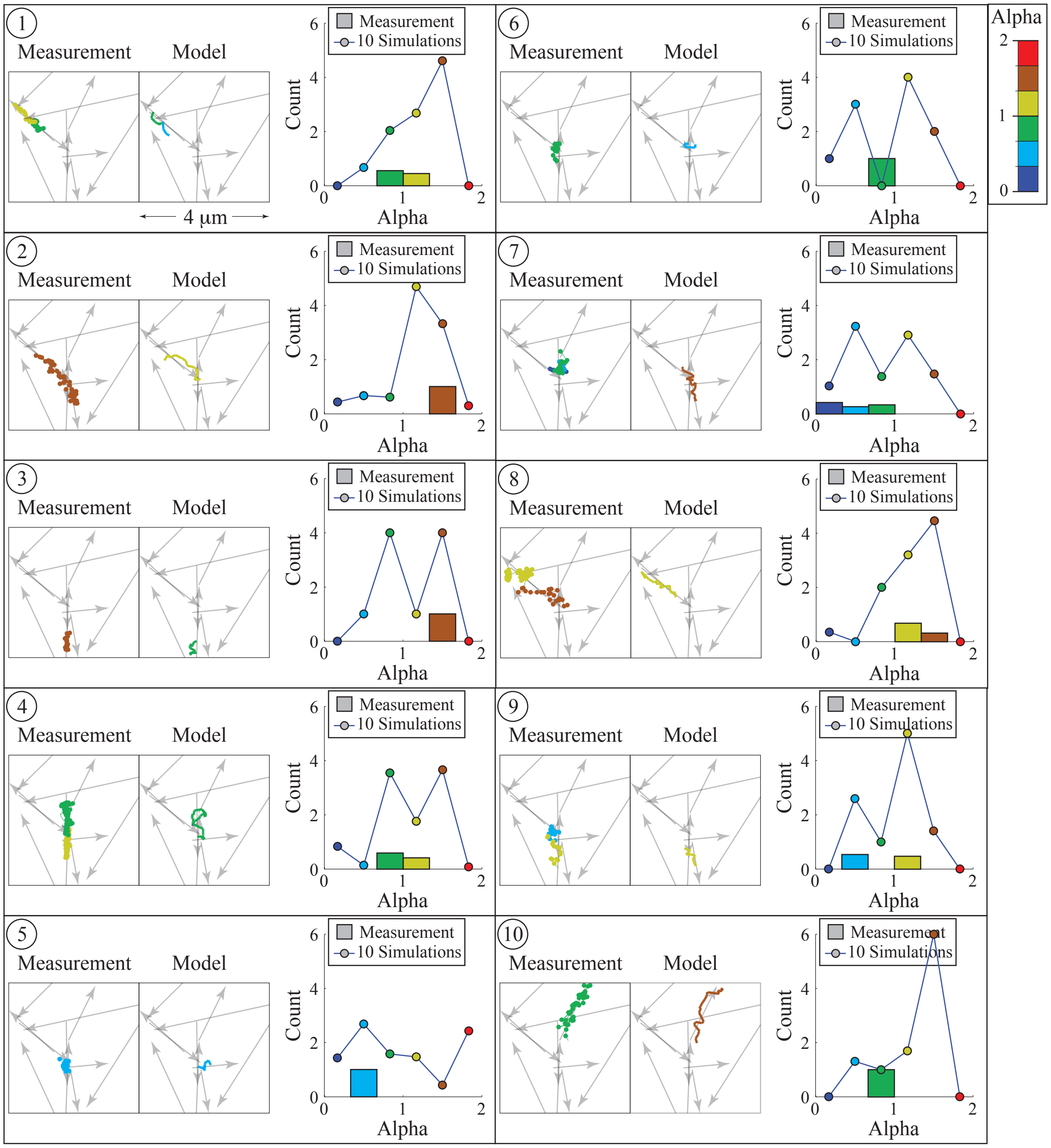


Figure S5: Simulation results and measurements from the unbranched network. Each panel, labeled 1-10, corresponds to a different measured trajectory observed on a $4\mu m \times 4\mu m \times 0.4\mu m$ section of the unbranched network. Inside each panel are, from left to right, i) the measured trajectory, ii) one of ten simulations, selected at random, and iii) a histogram of alpha value for the measurement and for the ten simulations. In each plot, color indicates alpha value, with warm colors indicating large (directed), and cold colors indicating small (stationary) values (scale at upper right).

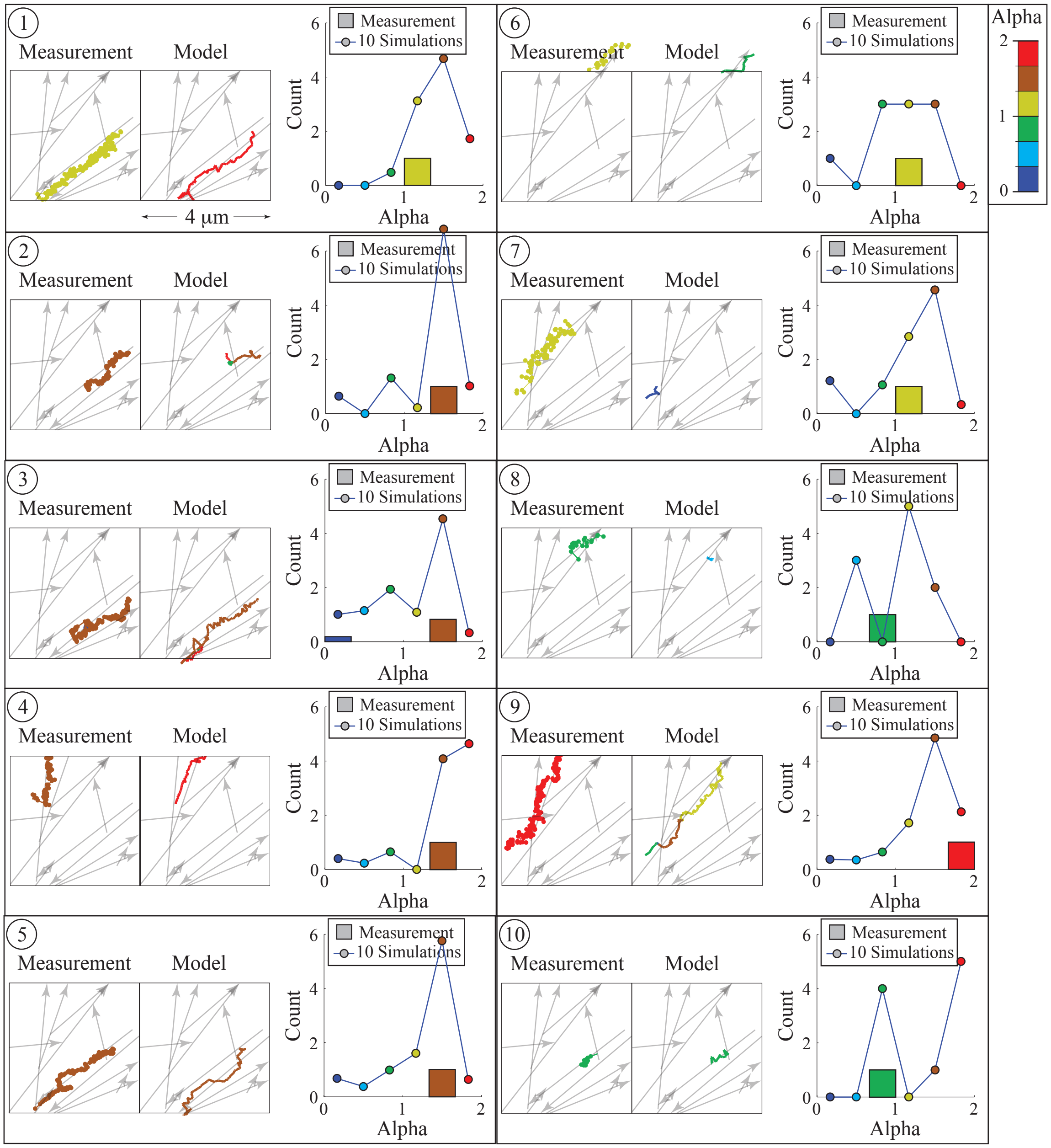


Figure S6: Simulation results and measurements from the Arp2/3-branched network. Each panel, labeled 1-10, corresponds to a different measured trajectory observed on a $4\mu$m $\times$ $4\mu$m $\times$ $0.4\mu$m section of the Arp2/3-branched network. Inside each panel are, from left to right, i) the measured trajectory, ii) one of ten simulations, selected at random, and iii) a histogram of alpha value for the measurement and for the ten simulations. In each plot, color indicates alpha value, with warm colors indicating large (directed), and cold colors indicating small (stationary) values (scale at upper right).

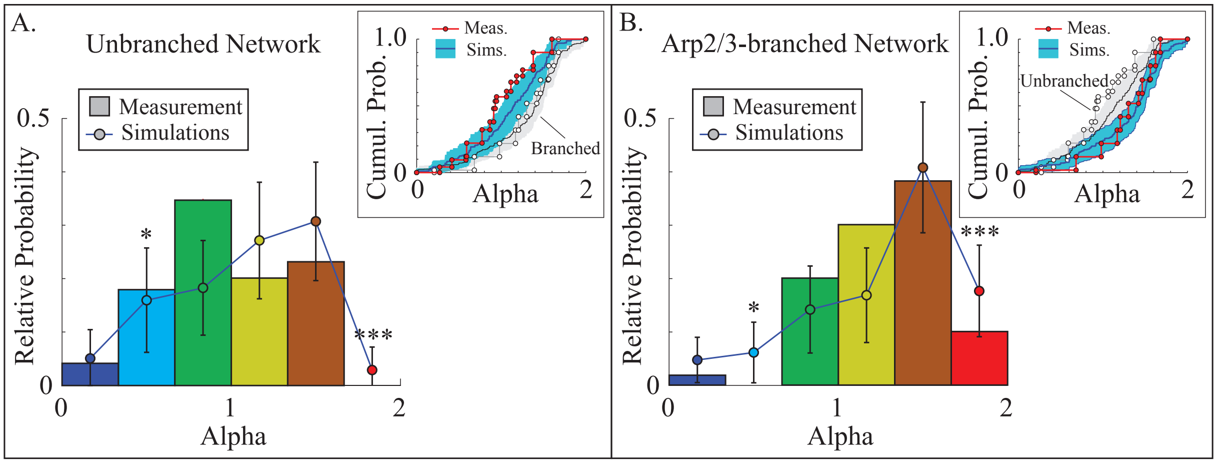


Figure S7: Model reproduces observed differences in mode of liposome motion between the two networks. In each plot, the filled histograms are the 10 measurements observed on the $4\mu m \times 4\mu m \times 0.4\mu m$ section of the unbranched (a) and Arp2/3-branched (b) networks. Mean of simulations of these measurements are indicated by the filled circles and error bars show standard deviation. Inset shows cumulative probability distributions, demonstrating that the differences do not depend on our choice of bin width in the histograms. $*$ indicates $p < 0.05$, $***$ indicates $p < 0.001$ (t-test).

##### 5.3 Analyzing the results

For a more quantitative comparison between the simulations and our experimental measurements, we generated histograms of alpha values for the simulated and measured trajectories on the two chosen local networks (Fig. S7). The histograms for the 10 experimentally observed trajectories in the $\sim 4\mu m \times 4\mu m \times 0.4\mu m$ regions (Fig. S7) show the same trends as the measured histograms for all $\sim 300$ measured trajectories (Fig. 3B,C main text). In particular, the Arp2/3-branched network has more directed and fewer stationary modes of motion.

To quantitatively compare the simulation results to our experimental observations, we needed to estimate both the expected histograms of alpha values from the simulations and also the uncertainty in those histograms, i.e. the mean and standard deviation for the height of each bin in the histogram. We estimated these quantities as follows. There are $10^{10}$ different combinations of 10 simulated trajectories with initial conditions matching the 10 measurements. We randomly picked 100 of these, and generated histograms for each. The mean and standard deviations of those 100 histograms are shown for both networks in Fig. S7. On both networks, the model is not significantly different from the experimentally observed measurements ($p > 0.05$, $\chi^2$-test). More importantly, there are significantly more directed trajectories ($p < 0.001$, t-test) and significantly fewer stationary trajectories ($p < 0.05$, t-test) on the Arp2/3-branched network, consistent with our observations. Thus, the model captures the differences in the modes of motion of liposome transport observed between the unbranched and the Arp2/3-branched networks.

### 6 Local geometry dictates type of liposome motion

The model successfully predicts observed differences between the Arp2/3-branched and unbranched networks. Given that the only input into the model is the network geometry, it seems likely that these differences arise from geometry. To further test this hypothesis, and to provide a more detailed understanding of how, precisely, geometry might dictate type of liposome motion, we performed a second series of simulations.

Within the *in vitro* unbranched networks, we selected 10 different local geometries or actin "motifs", defined by $\sim 2\mu\text{m} \times 2\mu\text{m} \times 0.4\mu\text{m}$ network regions. Five of these actin motifs were associated with nine different motile trajectories ($\alpha = 1.428 \pm 0.129$), and were therefore termed "motile actin motifs." The other five motifs were termed "non-motile actin motifs," since they were associated with nine different stationary trajectories ($\alpha = 0.738 \pm 0.309$). We then ran 50 simulations on each motif. In order to ensure some variability in initial conditions, both in terms of the number and position of attached myosin motors, and the position of the liposome azimuthally on the actin filament, each simulation was run as follows. The liposome was positioned on the same actin filament that the experimentally observed trajectory began, but with the initial position varied by starting closer to the minus end of the actin filament. Once the simulated trajectory was complete, we determined the starting point by finding the time at which the simulated trajectory reached the experimentally observed liposome's starting position. In some cases, particularly in dense actin networks, the simulated liposome never reached the observed liposome's starting point (e.g., because it switched to another actin filament prior to reaching the observed liposome's starting point), in which case we did not analyze the simulated trajectory.

We analyzed these simulated trajectories in the same way as we analyzed the *in vitro* experiments. The model reasonably reproduces the experimental measurements (Fig. S8). Further, on the motile actin motifs, we observed significantly more directed trajectories ($p < 0.001$, t-test) and significantly fewer stationary trajectories ($p < 0.001$, t-test) than on the non-motile actin motifs (Fig. S9). These results add further support to the idea that the differences we observe between trajectories arise from actin geometries on the scale of a few microns. More importantly, however, we can now determine the underlying cause for these differences.

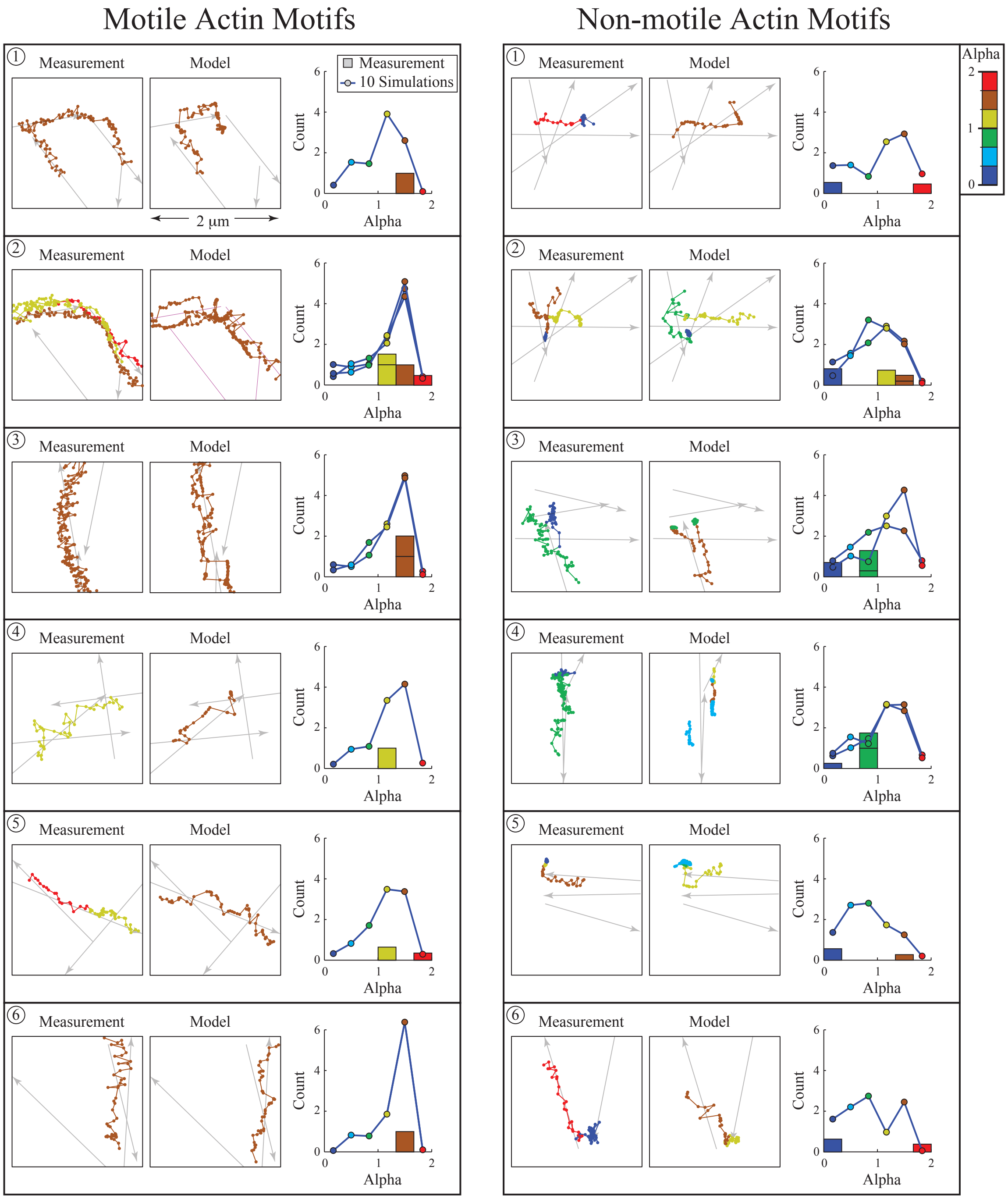


Figure S8: Model reproduces mode of motion on local geometries within the unbranched network. Left shows motile actin motifs, five local geometries associated with nine motile trajectories ($\alpha = 1.428 \pm 0.129$); right shows non-motile actin motifs, five local geometries associated with stationary trajectories ($\alpha = 0.738 \pm 0.309$). Note that the geometries in panel 1 and 2 are identical, but the trajectories start on different actin filaments. Each panel shows, from left to right, i) measured trajectory(s), ii) the simulated trajectory(s) closest (in least squares sense) to the measured trajectory(s), iii) a histogram of alpha value for the measurement(s) and for the $\sim 50$ simulations, scaled to be comparable to the histograms in Figs. S4 and S5.

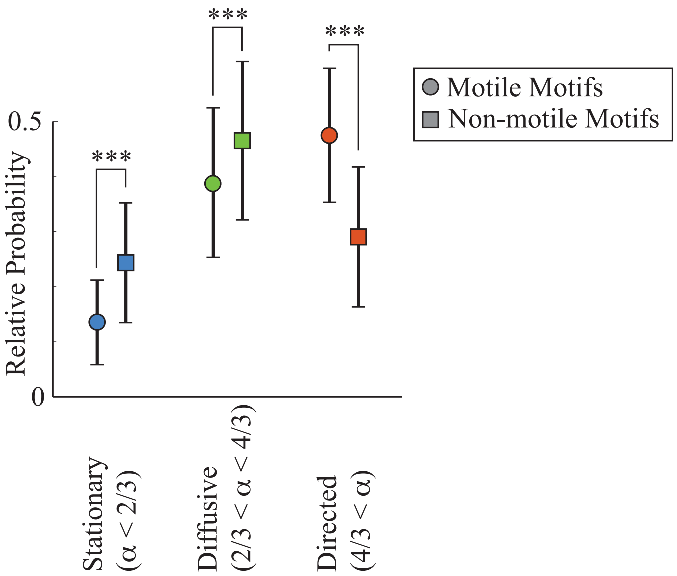


Figure S9: Simulations show more directed and less stationary trajectories on the motile, compared to the non-motile, actin motifs. Alpha values from nine simulated trajectories on the motile (and non-motile) actin motifs were calculated, and binned into the three categories, stationary ($\alpha < 2/3$), diffusive ($2/3 < \alpha < 4/3$) and directed ($4/3 < \alpha$). Symbols show mean values from simulations of the measurements, error bars show standard deviation. $***$ indicates $p < 0.001$ (t-test).

#### 6.1 Differences in mode of liposome motion arise from pauses in the trajectories

The majority of the experimentally measured trajectories on the non-motile actin motifs showed extended "pauses," where the liposome remained approximately motionless for an extended period of time (> 1s). The experimentally measured trajectories on the motile actin motifs did not show similar pauses. To quantify this effect, we plotted the distance between the liposome's current and initial position as a function of time ($d(t)$). We then fit the resulting data with a piecewise linear, and piecewise constant function of the form

$$d(t) = \begin{cases} at + b & : \quad t < t_1 \\ at_1 + b & : \quad t_1 \leq t < (t_1 + t_{pause}) \\ at_1 + b + c(t - t_1 - t_{pause}) & : \quad (t_1 + t_{pause}) \leq t \end{cases} \tag{2}$$

These best fits, along with the measured $d(t)$, the actin motifs and trajectories to which they correspond are shown in Fig. S10.

The experimentally observed trajectories on the non-motile actin motifs showed longer pauses ($t_{pause} = 1.66 \pm 1.43$s) than those on the motile actin motifs ($t_{pause} = 0.23 \pm 0.22$s, significantly different $p < 0.01$ t-test). As one might expect, these pauses were associated with low $\alpha$ values (Fig. S11). The fraction of time spent paused ($t_{pause}/t_{tot}$, where $t_{tot}$ is the total trajectory time), is negatively correlated with $\alpha$ (with a slope of $-1.14$, $p < 0.001$ F-test). This result shows that pauses and low $\alpha$ values generally occur together. To show that the pauses cause the low $\alpha$ values, we calculated the average $\alpha$ value for the trajectories on the non-motile actin motifs during the times that the liposome was not paused. The resulting $\alpha$ ($\alpha = 1.20 \pm 0.45$) is not significantly different ($p > 0.05$, t-test) from the $\alpha$ value for the trajectories on the motile actin motifs ($\alpha = 1.43 \pm 0.13$). Therefore, the significant differences we observe between the experimentally measured liposome trajectories on motile and non-motile actin motifs are a result of the pauses (Fig. S11).

The picture that emerges from this analysis is that myosin Va motors transport liposomes along an actin track until they encounter a particular actin geometry, where they pause. The length of these pauses dictates the efficiency of transport. Since our model captures the gross behavior of our experiments,

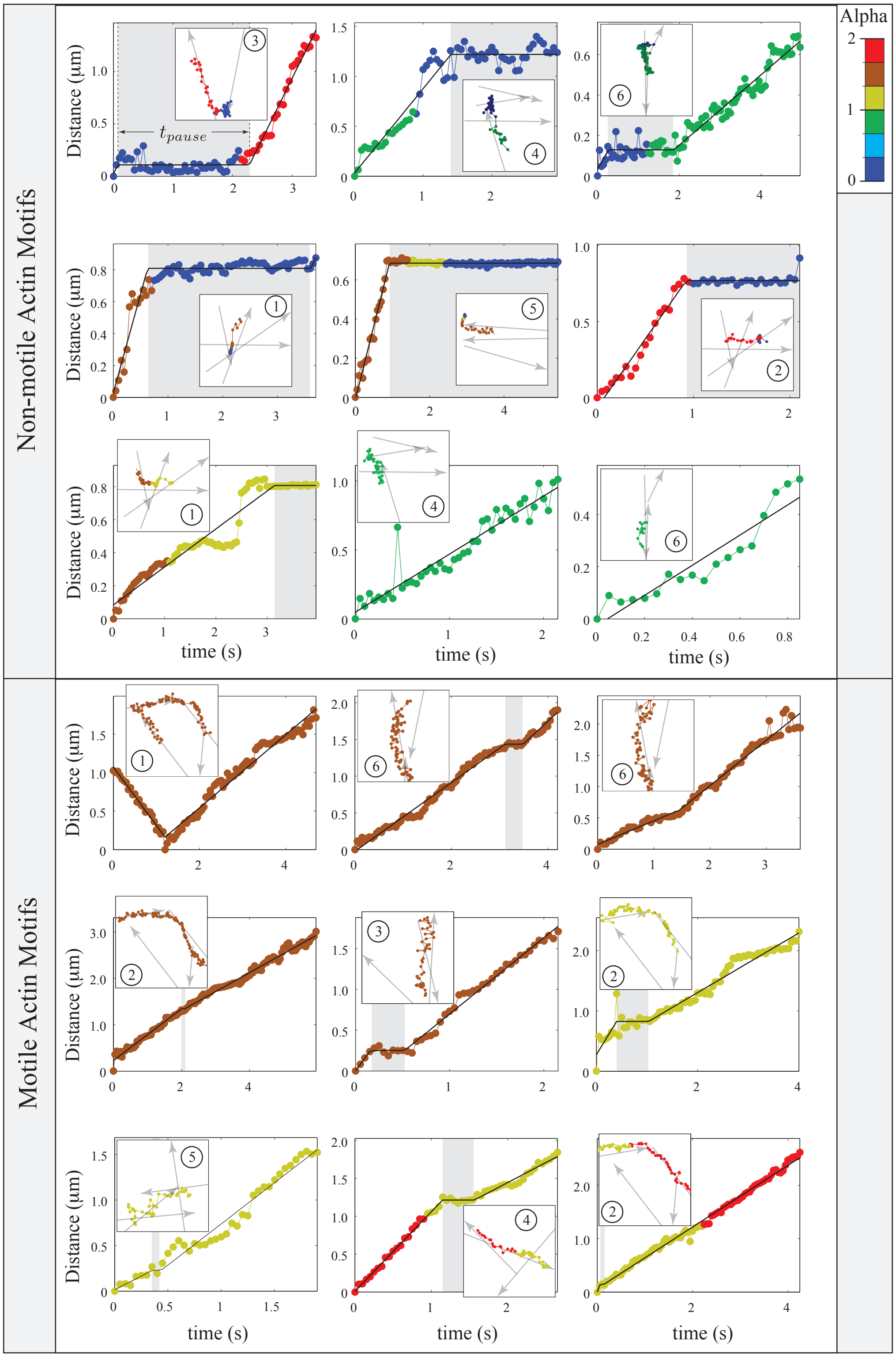


Figure S10: Measured trajectories on non-motile actin motifs (top) show distinct > 1s pauses (shaded in gray) that do not occur on motile actin motifs (bottom). Each plot shows distance from the initial liposome position as a function of time. The solid line is a piecewise linear/constant fit (Eq. 2). Inset shows the measured trajectory and geometry; the number in the upper right corresponds to the numbers given in Fig. S8. In each plot, color indicates alpha value, with warm colors indicating large (directed), and cold colors indicating small (stationary) values (scale at upper right).

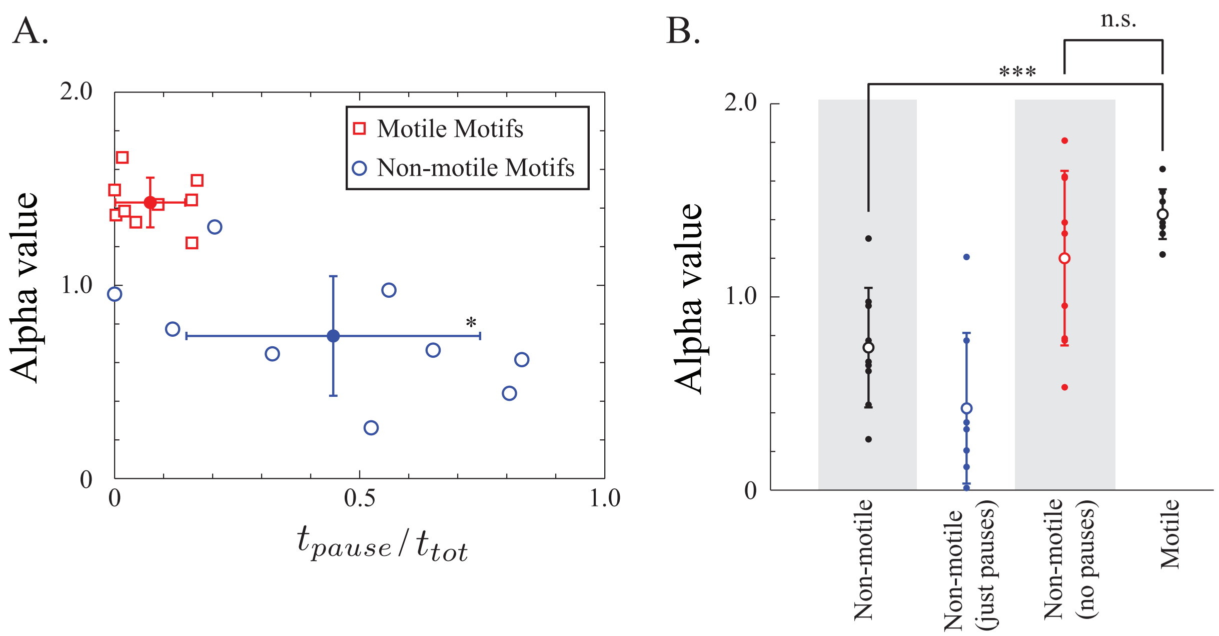


Figure S11: The low alpha values of experimentally measured trajectories on non-motile actin motifs are due to pauses. A. Alpha value negatively correlates with the proportion of the total trajectory time ($t_{tot}$) spent in a pause ($t_{pause}$). Trajectories on non-motile actin motifs pause for a longer time and have lower alpha values than those on motile actin motifs. B. If the pauses are removed, trajectories on non-motile actin motifs do not have different alpha values than those on motile actin motifs. $*$ indicates $p < 0.05$ (t-test); $***$ indicates $p < 0.001$ (t-test).

correctly predicting which actin geometries are associated with more or less efficient transport (Figs. S7 and S9), we used the model to understand the basis for these pauses. However, in order to do so, we first ensured that the model generates pauses at approximately the same spatial location as in our experimental observations.

#### 6.2 Model pauses at the same locations as our measurements

To determine whether the model pauses at approximately the same spatial location as in our experimental observations, we considered the six stationary trajectories that showed extended pauses ($>$ 1s, Fig. S12). We then examined the simulations run on the same actin motifs, looking for pauses. In all six cases, we identified at least one trajectory that paused close to (within $\sim$ 300nm) the same location as our measurements. Like in the experiments, in the simulations these pauses were associated with periods of low alpha ($\alpha < 0.667$). Therefore, it seems that the model is identifying the same geometries that we observe experimentally, so that we can look to the model to understand why these pauses occur.

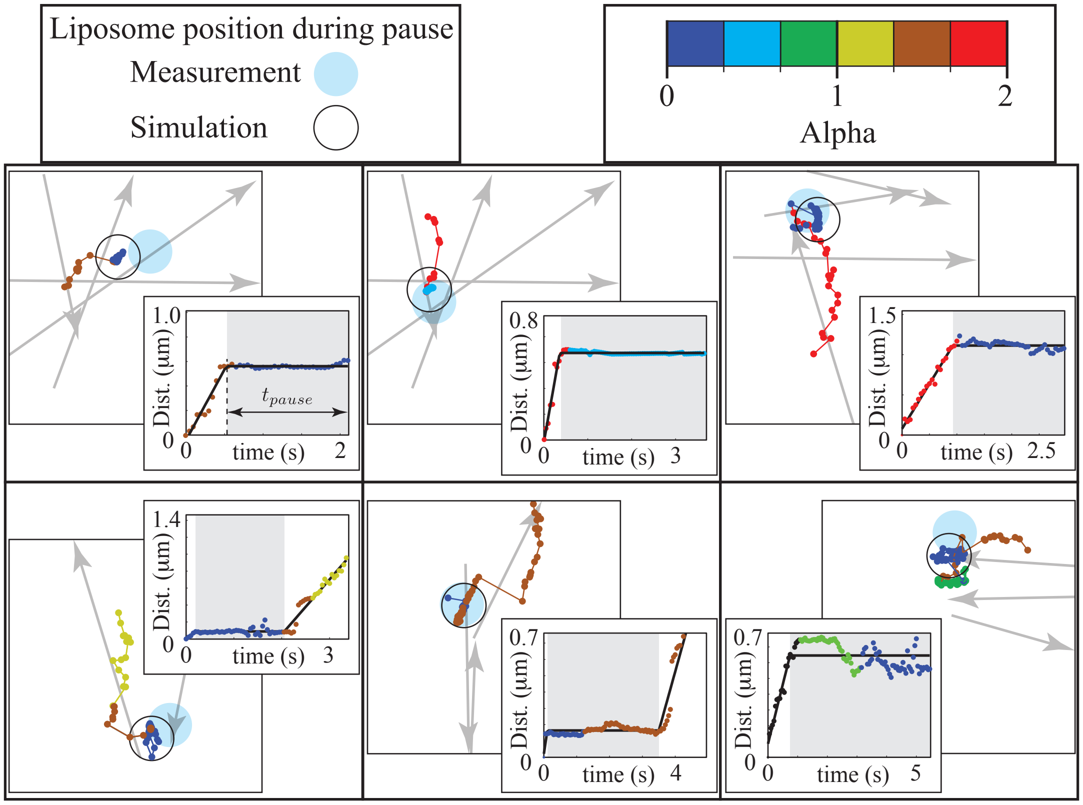


Figure S12: Simulations on non-motile actin motifs show pauses in the same 3D location as the measurements. For each non-motile actin motif, at least one of the ∼ 50 simulations paused in the same (i.e. within ∼ 300nm) 3D location as the measured pause. To give scale, the liposome is shown actual size (350nm). Insets show position vs. time plots, c.f. Fig. S10. In each plot, color indicates alpha value, with warm colors indicating large (directed), and cold colors indicating small (stationary) values (scale at upper right).

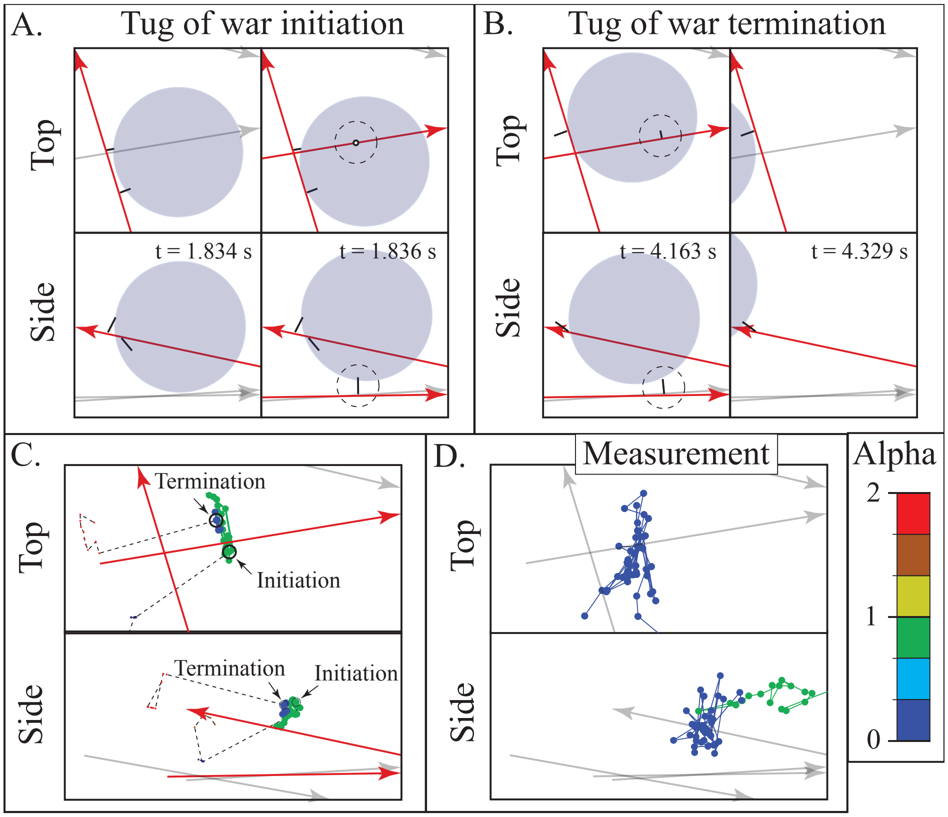


Figure S13: In the model, pauses occur due to tug of wars between myosin motor teams interacting with different actin filaments. In each panel, The top row shows a top view, the bottom row a side view. For panels A-C, actin filaments with one or more myosin bound are shown in red, and with no myosin bound are shown in gray. A. The liposome position, motor orientation, and actin position just prior to, and just after the initiation of the tug of war. B. The liposome position, motor orientation, and actin position just prior to, and just after the termination of the tug of war. C. A $1\mu m \times 0.5\mu m$ region with the simulated trajectory, along with with the location of tug of war initiation and termination, and the liposome position during the tug of war. The liposome position before and after the tug of war is shown as a dashed line. D. The measured trajectory, at the same spatial position as panel C. In plots C. and D., color indicates alpha value, with warm colors indicating large (directed), and cold colors indicating small (stationary) values (scale at lower right). A movie of this simulation can be seen in Movie S8.

##### 6.3 Pauses occur due to tug-of-wars

We examined three of the non-motile actin motifs in more detail, in an attempt to identify (1) whether the pauses in the model were due to a tug of war, and (2) if so, which actin filaments were involved. In all cases, the initiation of a tug of war was correlated with a long (1s or longer) pause and a period of low $\alpha$ (Figs. S13-S15 and Movies S8-S10). The termination of this tug of war ended the pause and the period of low $\alpha$. All tug of wars occurred between motor teams interacting with two actin filaments. In two cases, the liposome continued along the original filament; in one case the liposome switched filaments at the termination of the tug of war. Interestingly, all three of the actin filament pairs at which the pauses and tug of wars were observed formed an intersection angle of greater than 90 degrees.

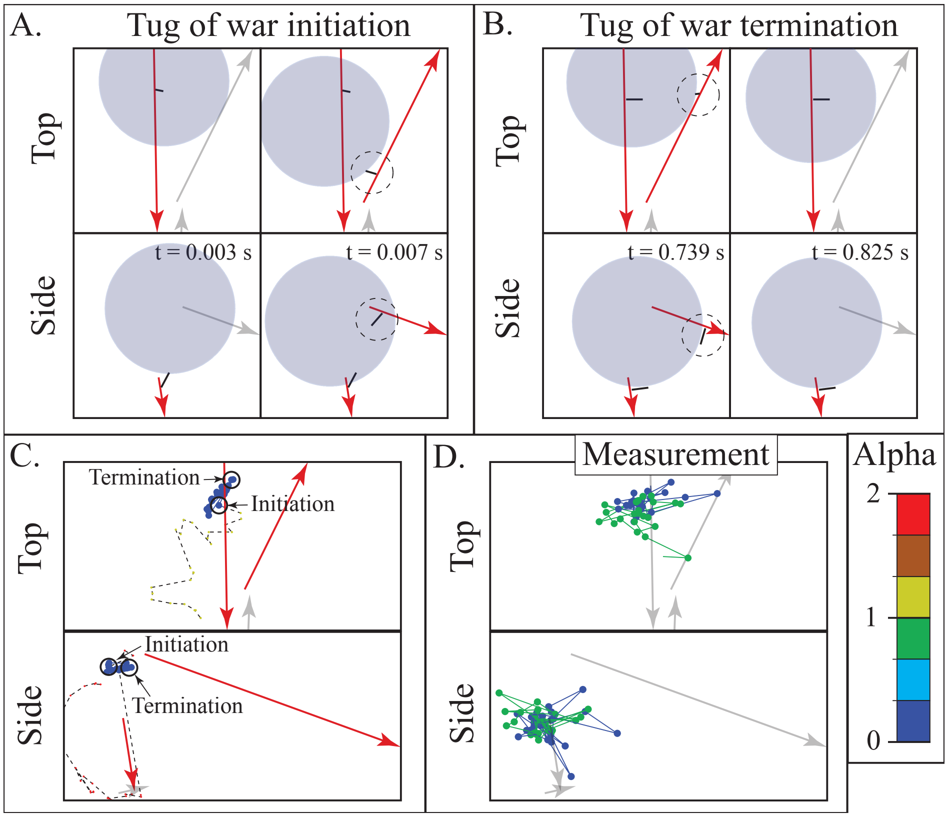


Figure S14: See caption for Fig. S13. A movie of this simulation can be seen in Movie S9.

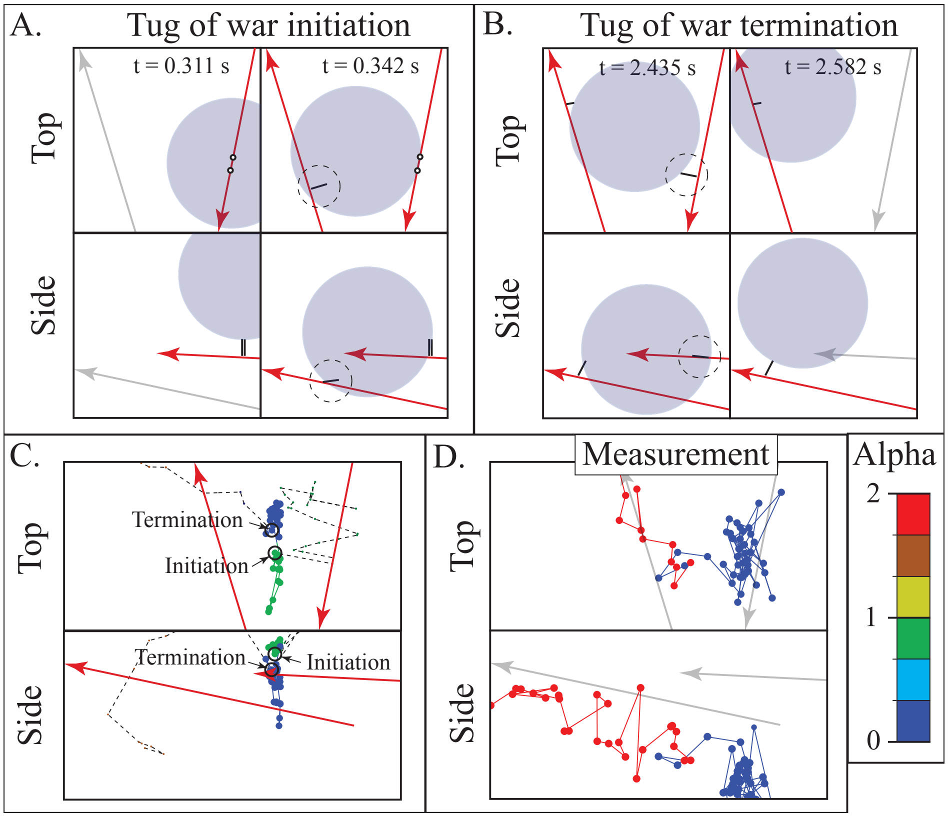


Figure S15: See caption for Fig. S13. A movie of this simulation can be seen in Movie S10.

#### 6.4 Force dependent detachment/stepping of myosin explains model dependence on local geometry

Given that pauses are caused by tug of wars between motor teams, and given that tug of wars generally occur between motor teams interacting with a pair of actin filaments, understanding how motor teams navigate actin intersections will give us an understanding of how and why tug of war duration depends on local geometry. We therefore performed a series of simulations at intersection angles ranging from $\theta = 33.75°$ to $\theta = 146.25°$. These bounds were picked because angles outside this range (i.e. approximately parallel or antiparallel actin filaments) do not have a discrete intersection point, where a single tug of war can occur, but rather a long intersection zone, through which multiple tug of wars can occur. Within this range of angles, and besides the end points ($\theta = 33.75$ and $\theta = 146.25°$), we ran simulations at $\theta = 45, 90$ and 135°.

In addition to intersection angle, we expected that the separation distance between filaments would influence the tug of wars between motor teams. We therefore examined filament separations of $0, 100, 200, 300$ and 400nm for each intersection angle (a total of 25 different conditions). For each different condition, we ran 50 simulations starting with a single motor bound to the actin filament on which it will approach the intersection and at a distance between 745 and 800nm from the intersection. Within the 55nm starting zone, the motor was initially bound to a random actin monomer, which (1) effectively randomizes the initial condition, and (2) ensures that the system is approximately in steady-state by the time the liposome reaches the intersection. We re-ran any simulations that failed to reach the intersection. Simulations were terminated either when all the motors detached (after reaching the intersection), or the liposome passed 500 nm beyond the intersection.

After running the simulations, we identified pauses by fitting a curve of the distance from the liposome's center of mass to the intersection $d(t)$ as a function of time with a piecewise linear/constant function (Eq. 2). The resulting cumulative probability distribution of pause times ($P(t_{pause})$) was then fit with a double exponential.

$$P(t) = 1 - \left(f_{slow}(z)e^{-t/t_{slow}(\theta)} + (1 - f_{slow}(z))e^{-t/t_{fast}}\right) \quad (3)$$

We used this function because we expected that sometimes the motors on the liposome surface would interact with the crossing actin filament, and sometimes they would not. In the former case, occurring with probability $f_{slow}$, we would expect to see longer pauses (of average duration $t_{slow}$) than in the latter case (where there would be no pause, but random fluctuations in liposome position would likely result in best fit curves with a short pause, $t_{fast}$). While the underlying process that governs the lifetime of a tug of war is likely not described by a single exponential, we nevertheless chose this function because we expected that such a distribution would capture the gross changes in tug of war lifetime we observed in our simulations. Interaction probability, $f_{slow}$, was anticipated to be a function of actin filament separation (since tug of wars should happen only rarely for the largest intersection separation distance of 400 nm) and the tug of war time $t_{slow}$ was anticipated to be a function of intersection angle $\theta$. The remaining parameter, the average lifetime of the short pauses ($t_{fast}$), was expected to remain constant, independent of $\theta$ and $z$. Consistent with these expectations, using a value of $t_{fast} = 0.667$s, and tuning the parameters $f_{slow}$ and $t_{fast}$ to fit the cumulative probability distribution of $t_{pause}$ from the simulations (using Matlab's optimization function `fminsearch` to minimize the sum of the squared difference between Eq. 3), Eq. 3 gave reasonable agreement (see Fig. S16A) and allowed us to estimate $f_{slow}$ and $t_{slow}$ for all 25 simulation conditions.

We observe a $\theta$ dependence in pause lifetime ($t_{slow}$), and no obvious $z$ dependence; conversely, we observe a $z$ dependence in pause probability $f_{slow}$, and no obvious $\theta$ dependence (Fig. S16B,C). Supposing that this $\theta$ dependence arises from the force dependence of the motors (which have force-dependent reaction

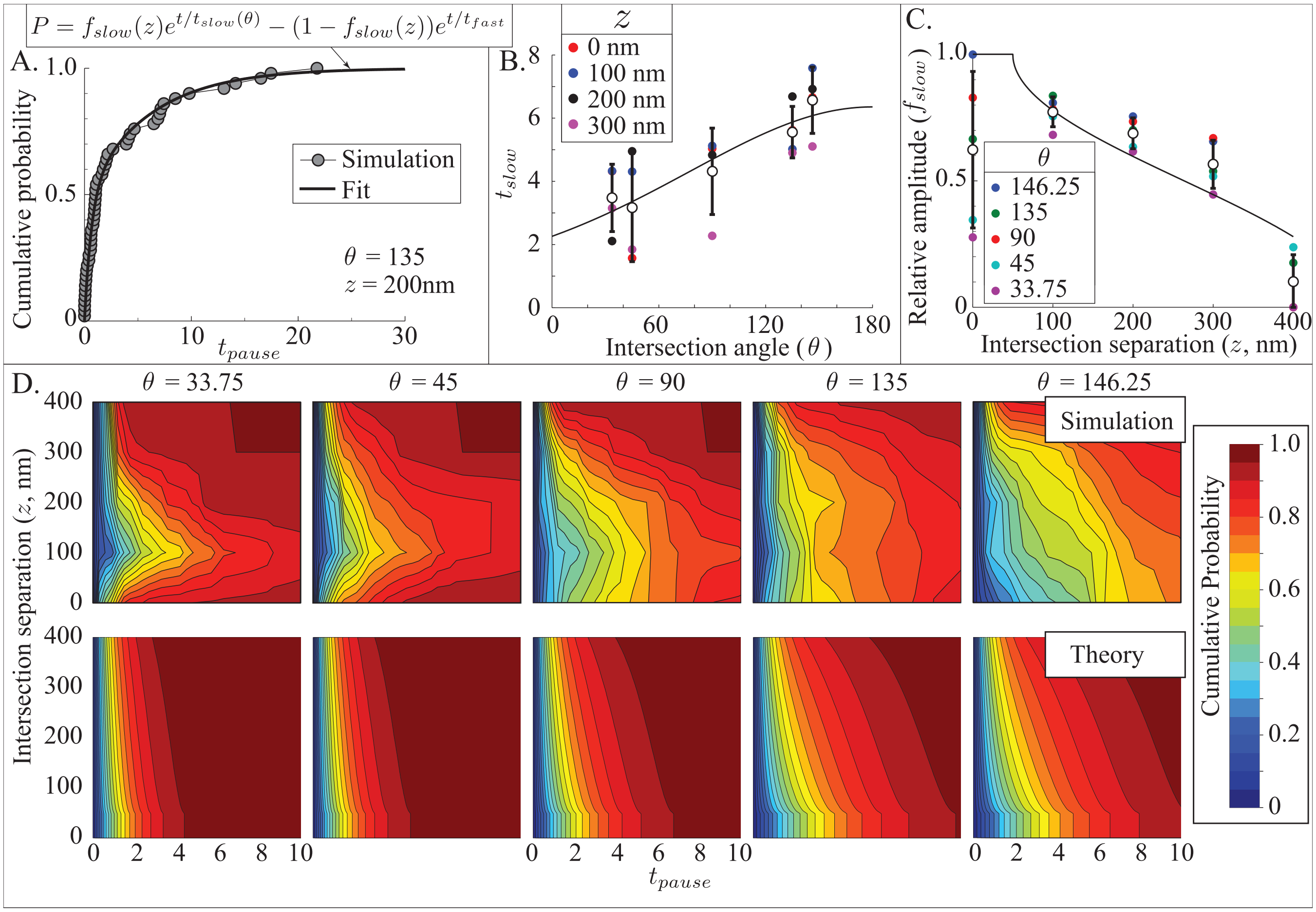


Figure S16: Force-dependent detachment/stepping of MyoVa predicts the duration of liposome pauses at actin filament intersections. A. Results of simulations at one intersection (angle between filaments, $\theta = 135°$; separation between filaments, $z = 200$nm). The cumulative probability of liposome pause duration is described by a double exponential, allowing the determination of pause probability ($f_{slow}$) and average duration ($t_{slow}$). B. Average pause duration, $t_{slow}$, is independent of $z$, and reasonably described by Bell's approximation, Eq. 4 (solid line)), a description of force-dependent chemical reactions. C. Pause probability is described by the probability that the liposome contacts the crossing filament (Eq. 6, solid line). D. Combining force dependent chemistry (Eq. 4) and interaction probability (Eq. 6) predicts pause duration at any actin intersection of angle $\theta$ and separation $z$ (top row, simulation results; bottom row, prediction). Note: $z = 400$nm is not included in the plot of $t_{slow}(\theta)$ because the liposome almost never interacted with the crossing filament at such a large separation.

rate that follow Bell's approximation), then we would expect to observe a functional form like

$$t_{slow} = t_0 \exp(c \cos(\frac{180^\circ - \theta}{2})) \tag{4}$$

where $t_0$ is a constant representing tug of war time in the absence of intermolecular forces and $c$ is a constant that is roughly the forces between motor teams $F$, times the force-dependent distance $\lambda$ from Bell's approximation divided by Botlzmann's constant times temperature, $k_B T$. The $\cos(\frac{180^\circ - \theta}{2})$ arises because the reaction rates depend on the component of force along the actin filament. Best-fit parameter values (from minimizing sum of the squared difference between Eq. 4 and mean $t_{slow}$ using Matlab's `fminsearch` function, fit shown in Fig. S16B) are $t_0 = 2.258$ s, and $c = 1.035$.

We observe a $z$ dependence in interaction probability, $f_{slow}$. At the largest filament separations ($z =$ 400nm), the amplitude was lowest (nearly 0) and generally increased with decreasing $z$. Given an actin intersection with separation $z$ and an azimuthal approach angle $\phi$, we have previously proposed (Lombardo et al. 2017) that motors on the liposome can interact with a crossing actin filament if

$$z < R_{eff} \cos(\phi) + \ell + R_{eff} \tag{5}$$

where $R_{eff}$ is the effective radius of the liposome and $\ell$ is the length of the myosin Va molecule. Supposing that azimuthal approach angle is uniformly distributed between 0 and 360°, this suggests an interaction probability of

$$f_{slow} = \begin{cases} 1 & : \quad z < \ell \\ \text{acos}\left(\frac{z - \ell - R_{eff}}{R_{eff}}\right)\frac{1}{\pi} & : \quad \ell < z < 2R_{eff} + \ell \\ 0 & : \quad 2R_{eff} + \ell < z \end{cases} \tag{6}$$

Note that using $R_{eff} = R$ in Eqs. 5 and 6 assumes that the liposome is in contact with the actin filament on which it travels (Lombardo et al. 2017); however, it is possible that the liposome is anywhere between $R < R_{eff} < R + \ell$ away from the actin filament on which it travels. Fitting this equation to our simulations gave an estimate of $R_{eff} = 213$nm (Fig. S16C).

Together, Eqs. 4 and 5 allow us to predict pause duration at any actin intersection of angle $\theta$ and separation $z$. This simplified description does a reasonable job recapitulating the simulation results (Fig. S16D). Importantly, the agreement between the simple description and the simulations suggests that the theory reasonably describes (1) whether a tug of war will happen and (2) how long that tug of war will last. Further, in the theory, tug of wars occur when the liposome interacts with a crossing filament, and the tug of wars last longer at large intersection angles because then the motors apply forces on each other that slow their detachment, thereby delaying the resolution of the tug of war.

##### 6.5 Liposome motion in the two networks is consistent with differences in local geometry

Given the analysis so far, we predict that the differences we observe between the Arp2/3-branched and the unbranched networks arise because, in the Arp2/3-branched network, some of the random actin intersections are replaced by Arp2/3 branches. Given a branch angle of 70°, we would predict that the motor teams can navigate relatively efficiently through these geometries (see Fig. S16). Thus, in general, transport is more directed on the Arp2/3-branched networks, in agreement with our observations and simulations.

To quantitatively test this idea, we ran 50 simulations through an Arp23 geometry. From these simulations, we generated a histogram of alpha values, normalized so that the sum of the height of all bins is 1. We then generated a series of histograms, formed by combining a fraction (say, $f$) of these histograms with a fraction $(1 - f)$ of histograms simulated on the unbranched network. Given that the alpha values from the unbranched network represent the result of liposomes passing through random intersections, then each combined histogram represents the results of liposomes passing through a network composed of a mixture

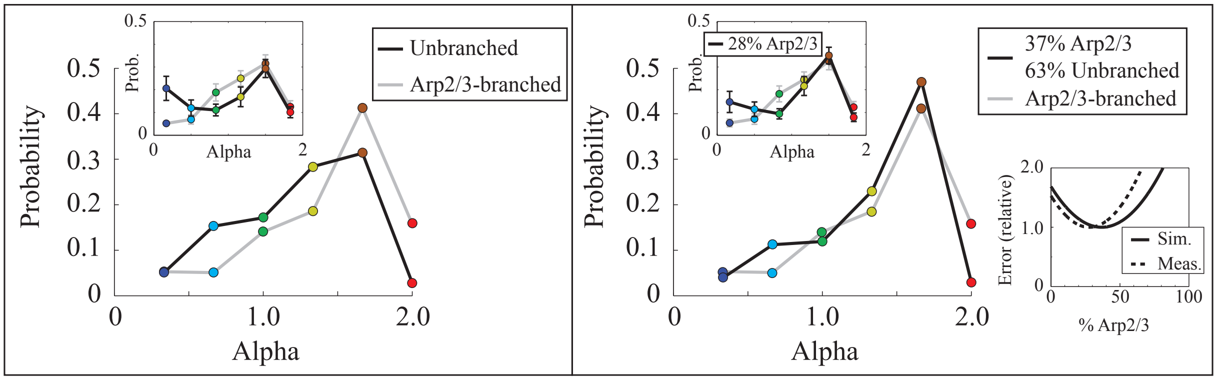


Figure S17: Left, original histograms of simulations on Arp2/3-branched and unbranched networks (measurements in inset). Right, the simulations on the Arp2/3-branched network are well-approximated by adding simulations through Arp2/3 branches to the simulations on the unbranched network in proportions of 37% and 63%, respectively (measurements in upper inset). Lower right inset, dashed line, shows mean squared difference between histograms from the Arp2/3-branched network and histograms from combining different fractions of histograms from simulations through Arp2/3 intersections and histograms from simulations on the unbranched network. The minimum error occurs at 37% Arp2/3, the value used to generate the histograms at right. Solid line shows similar results when the experimentally measured histograms one the unbranched and Arp2/3-branched networks are used instead of the simulated histograms. Note that the mean squared error is scaled to be 1 at the minimum.

of random and Arp2/3 intersections with relative probabilities $1 - f$ and $f$, respectively. We assessed the mean squared difference between the alpha value histogram from simulations on the Arp2/3-branched network, and histograms generated by combining different fractions of random and Arp2/3 intersections. This mean squared difference is minimized with 37% ($f = 0.37$) Arp2/3 branches and 63% ($1 - f = 0.63$) random intersections (Fig. S17, lower right inset). The agreement between the two histograms is reasonable (Fig. S17). Additionally, the same procedure works for the experimental measurements as well (Fig. S17, upper left inset). Furthermore, the proportion of Arp2/3 intersections is broadly consistent with the proportion of Arp2/3 intersections predicted from our measurements of Local Polarity Alignment ($\sim 15\%$, Fig. 5C, main text). Overall, then, these results support our idea that the increase in directed motion observed on the Arp2/3-branched network arises from the relatively efficient way that myosin Va motors navigate a liposome through an Arp2/3 branch.

**Supplemental Information Appendix:**

**Supplemental Movie S1**: The polarity of actin filaments in a 3D network determined by reporter cargos. The trajectories of fluorescent single myosin reporter cargos (magenta) are overlaid on a 3D STORM reconstruction of Alexa-647-phalloidin-labeled actin network (3D STORM color scale shown in Fig. 1B of the main text). Single myosin motors move towards the plus-end of the actin filament to which they attach. Thus, tracking the movement of multiple reporter cargos (white) specifies the polarity of each actin filament in the 3D network. This movie was created by combining segments of a continuous 4000 frame movie to show all filaments' polarity being identified. Movie playback 2x real time. Scale 500nm.

**Supplemental Movie S2:** Directed liposome motion is observed on aligned actin filament networks. Results of one simulation of a 350nm liposome (blue) transported by 10 freely diffusing MyoVa motors (yellow cylinders) along a network of four actin filaments (yellow arrows point toward the filaments' plus end). The position and polarity of the actin filaments are taken from our experimental measurements of an unbranched actin filament network. The motors quickly overcome multiple scenarios when the liposome is in proximity of two 'nearly polarity aligned' actin filaments to maintain directed transport. Note that only actin-bound MyoVa motors are pictured, and that a maximum of three motors can be simultaneously bound to actin (see Lombardo et al. 2017 for justification of this assumption). Movie playback real time.

**Supplemental Movie S3:** Movies of the directed motion data presented in Fig. 7A of the main text. Left: Overlay of fluorescent 350nm liposome with ~10 myoVa motors (magenta) on 3D STORM reconstruction of Alexa-647-phalloidin-labeled actin network (3D STORM color scale shown in Fig. 1B of the main text). The liposome is transported in a directed mode along the 'aligned/nearly aligned' actin network. Right: To-scale reconstruction of the 3D actin network shown at left (created as shown in Fig. 4 of the main text, and described in Methods) with arrow heads indicating the plus end of the actin filaments. The 350nm liposome (magenta) follows the tracked trajectory (red) of the experimentally collected data. A gray halo surrounds the liposome, as an estimate of where myoVa motors can reach. Movie playback is real time. Scale 500nm.

**Supplemental Movie S4:** Movies of the diffusive-like data presented in Fig. 7B of the main text. Left: Overlay of fluorescent 350nm liposome with ~10 myoVa motors (magenta) on 3D STORM reconstruction of Alexa-647-phalloidin-labeled actin network (3D STORM color scale shown in Fig. 1B of the main text). The liposome is transported in a diffusive-like mode along the 'poorly/nearly aligned' actin network. Right: To-scale reconstruction of the 3D actin network shown at left (created as shown in Fig. 4 of the main text and described in Methods) with arrow heads indicating the plus end of the actin filaments. The 350nm liposome (magenta) follows the tracked trajectory (red) of the experimentally collected data. A gray halo surrounds the liposome, as an estimate of where myoVa motors can reach. Movie playback is real time. Scale 500nm.

**Supplemental Movie S5:** Diffusive liposome motion is observed on poorly-aligned actin filament networks. Results of one simulation of a 350nm liposome (blue) transported by 10 freely diffusing MyoVa motors (yellow cylinders) along a network of four actin filaments (yellow arrows point toward the filaments' plus end). The position and polarity of the actin filaments are taken from our experimental measurements of an unbranched actin filament network. The cargo meanders between 'poorly aligned' filaments creating diffusive-like motion as motors engage in and quickly resolve multiple tug-of-wars. Note that only actin-bound MyoVa motors are pictured, and that a maximum of three motors can be simultaneously bound to actin (see Lombardo et al. 2017 for justification of this assumption). Movie playback real time.

**Supplemental Movie S6:** Movies of the stationary mode of motion data presented in Fig. 7C of the main text. Left: Overlay of fluorescent 350nm liposome with ~10 myoVa motors (magenta) on 3D STORM reconstruction of Alexa-647-phalloidin-labeled actin network (3D STORM color scale shown in Fig. 1B of the main text). The liposome experiences a stationary mode on the 'not aligned' actin network. Right: To-scale reconstruction of the 3D actin network shown at left (created as shown in Fig. 4 of the main text, and described in Methods) with arrow heads indicating the plus end of the actin filaments. The 350nm liposome (magenta) follows the tracked trajectory (red) of the experimentally collected data. A gray halo surrounds the liposome, as an estimate of where myoVa motors can reach. Movie playback is real time. Scale 500nm.

**Supplemental Movie S7:** Stationary liposomes are observed on not aligned actin filament networks. Results of one simulation of a 350nm liposome (blue) transported by 10 freely diffusing MyoVa motors (yellow cylinders) along a network of four actin filaments (yellow arrows point toward the filaments' plus end). The position and polarity of the actin filaments are taken from our experimental measurements of an unbranched actin filament network. The cargo becomes stationary between 'not aligned' filaments due to a tug-of-war between competing motors producing forces in opposing directions. Note that only actin-bound MyoVa motors are pictured, and that a maximum of three motors can be simultaneously bound to actin (see Lombardo et al. 2017 for justification of this assumption). Movie playback real time.

**Supplemental Movie S8:** Movie of the simulation shown in Fig. S13, demonstrating a pause due to a tug of war in the same 3D location as the experimental measurements show a stationary mode of motion. Small (50 nm) translucent blue spheres mark the 3D liposome position at which the stationary mode of motion was observed experimentally. 350 nm liposome (blue, with white 50 nm sphere marking the center of mass) transported by 10 freely diffusing MyoVa motors (yellow cylinders) along four actin filaments (yellow arrows point toward the filaments' plus end). During the tug of war, the position of the liposome's center of mass is marked by translucent 50 nm red spheres. The positions of those red spheres match the positions observed experimentally. Note that only actin-bound MyoVa motors are pictured, and that a maximum of three motors can be simultaneously bound to actin (see Lombardo et al. 2017 for justification of this assumption). Movie playback 0.5x real time.

**Supplemental movie S9:** Movie of the simulation shown in Fig. S14 of the supplement, demonstrating a pause due to a tug of war in the same 3D location as the experimental measurements show a stationary mode of motion. Small (50 nm) translucent blue spheres mark the 3D position at which the stationary mode of motion was observed experimentally. 350 nm liposome (blue, with white 50 nm sphere marking the center of mass) transported by 10 freely diffusing MyoVa motors (yellow cylinders) along three actin filaments (yellow arrows point toward the filaments' plus end). During the tug of war, the position of the liposome's center of mass is marked by translucent 50 nm red spheres. The positions of those red spheres match the positions observed experimentally. Note that only actin-bound MyoVa motors are pictured, and that a maximum of three motors can be simultaneously bound to actin (see Lombardo et al. 2017 for justification of this assumption). Movie playback 0.5x real time.

**Supplemental movie S10:** Movie of the simulation shown in Fig. S15 of the supplement, demonstrating a pause due to a tug of war in the same 3D location as the experimental measurements show a stationary mode of motion. Small (50 nm) translucent blue spheres mark the 3D position at which the stationary mode of motion was observed experimentally. 350 nm liposome (blue, with white 50 nm sphere marking the center of mass) transported by 10 freely diffusing MyoVa motors (yellow cylinders) along two actin filaments (yellow arrows point toward the filaments' plus end). During the tug of war, the position of the liposome's center of mass is marked by translucent 50 nm red spheres. The positions of those red spheres match the positions observed experimentally. Note that only actin-bound MyoVa motors are pictured, and that a maximum of three motors can be simultaneously bound to actin (see Lombardo et al. 2017 for justification of this assumption). Movie playback 0.5x real time.
